# Anatomically distinct OFC-PCC circuits relay choice from value space to action space

**DOI:** 10.1101/2020.09.01.277889

**Authors:** Maya Zhe Wang, Benjamin Y. Hayden, Sarah R. Heilbronner

## Abstract

Economic choice necessarily involves the transformation of abstract, object-based representations to concrete, action-based ones. This transformation is both determined and delimited by the neuroanatomical organization of the regions that implement it. In choice, the orbitofrontal cortex (OFC) plays a key role in both abstract valuation and cognitive mapping. However, determining the neural processes underlying this transformation has proven difficult. We hypothesized that difficulty stems from in part from the fact that the OFC consists of multiple functionally distinct zones that are distinguished by their differing contributions to the abstract-concrete transformation, and that these functions reflect their differing long-range projections. Here we identify two such subregions, defined by stronger or weaker bidirectional anatomical connectivity with the posterior cingulate cortex (PCC). We call these regions OFC*in* and OFC*out*, respectively. We find that OFC*in*, relative to OFC*out*, shows enhanced functional connectivity with PCC, as indicated by both spike-field coherence and mutual information. We find converging evidence that the OFC*in*-PCC circuit, but not the OFC*out*-PCC circuit, relays choice signals from an abstract value space to a concrete action space. Moreover, the OFC*in*-PCC circuit shows a putative bidirectional mutually excitatory pattern. Together, these results support the hypothesis that OFC-PCC subareal organization is critical for understanding the implementation of offer-action transformation in economic choice.

## INTRODUCTION

Among brain regions associated with economic choice, the orbitofrontal cortex (OFC) has attracted the lion’s share of attention (Bradfield & Hart, 2020; Kaplan et al., 2017; Padoa-Schioppa & Conen, 2017; Schoenbaum et al., 2009; Stalnaker et al., 2015; Wallis, 2007; Wikenheiser & Schoenbaum, 2016; Wilson et al., 2014; Rudebeck and Murray, 2014 and 2018). This region is associated with evaluation, value comparison, cognitive mapping, and prospection (Padoa-Schioppa, 2011; Rushworth et al., 2011; Schuck et al., 2016; Wallis, 2007; Wang et al., 2020; Wang & Hayden, 2017). Consequently, OFC is seen as playing a central role in choice. Furthermore, there is increasing attention being paid to functionally unique subdivisions of the OFC (Rudebeck & Murray, 2011 and 2018). For example, the medial OFC may be more associated with abstract valuation and learning processes (Noonan et al., 2010; Rushworth et al., 2011; Levy & Glimcher, 2014), whereas central OFC may help to associate stimuli with outcomes or signal outcome desirability (Niv, 2019; Wilson et al., 2014; Rudebeck et al., 2017), and the lateral OFC may signal resource availability (Rudebeck et al., 2017). However, these distinctions are based on coarse parcellation, and may not reflect the subtleties of anatomical and functional differentiation within this broad swath of cortex.

Economic choice requires the transformation of sensory and mnemonic information into actions (Cai & Padoa-Schioppa, 2014; Hare et al., 2011; Hayden & Moreno-Bote, 2018; Yim et al., 2019; Yoo et al., 2018). In other words, economic choice involves a transformation from an abstract (goods) space to a concrete (action) one (Padoa-Schioppa, 2011; Rangel et al., 2008). It is likely that the OFC plays a central role in this process. However, the nature of that role remains unclear. Many studies have emphasized the abstract side of OFC processing; however, a growing number of studies suggest that it may have an important spatial role as well (Yoo et al., 2018; Luk and Wallis, 2013; Strait et al., 2016; Roesch et al., 2006). The inconsistency across studies, along with the functional divisions explained above, raise the possibility that different parts of OFC may have heterogeneous functions. Defining, and working with, that heterogeneity may allow for more precise delineation of OFC function.

We hypothesized that the key to understanding the role of OFC in the transformation from abstract to concrete representations is through its connectivity with another region involved in economic choice: the posterior cingulate cortex (PCC). This region, located in the posteromedial cortex, has not received the same amount of scholarly scrutiny from decision neuroscientists as OFC. Nevertheless, the PCC has a confirmed spatial repertoire (Dean & Platt, 2006; Hayden et al., 2008; Olson et al., 1996; Spreng et al., 2009; Dean et al., 2004) and plays a central economic role (Barack et al., 2017; Hayden et al., 2008; Heilbronner & Platt, 2013; Kable & Glimcher, 2007; Pearson et al., 2009; Young & Mccoy, 2015). That is, while PCC has consistent responses to outcomes, those responses are spatially selective, perhaps due to the strong interactions between this region and the parietal cortex (Morecraft et al., 2004; Cavada et al., 1989; Pandya & Seltzer 1982). Finally, PCC has direct bidirectional communication with OFC (Kobayashi & Amaral, 2003; Morecraft et al., 2004; Pandya et al., 1981; Parvizi et al., 2006; Morecraft et al., 1992). We wanted to probe how the OFC-PCC circuit might facilitate transformations from abstract space to action space for choice.

## RESULTS

### OFC-PCC anatomical connectivity

We injected the tracer fluororuby in the PCC gyrus, centered at the border between area 23a and 30 (with some involvement of area 29, Paxinos et al., 2009). This injection resulted in widespread retrograde and anterograde labeling throughout the anterior and posterior cingulate cortices, parietal lobe (precuneus and intraparietal sulcus), medial temporal lobe (hippocampal formation), and frontal cortex (primarily dorsolateral prefrontal and orbitofrontal cortices). Projections to the OFC were particularly interesting for their specificity: cells and terminal fields were clustered around the medial orbital sulcus (mainly area 13a, but also including lateral 14O and caudal 11, based on Paxinos et al., 2009; **Figure 1A-C**). There were projections to other OFC subregions, but these were noticeably less dense. These results are consistent with other, similarly placed cases from the literature (Kobayashi & Amaral, 2003; Morecraft et al., 2004; Pandya et al., 1981; Parvizi et al., 2006; Morecraft et al., 1992). A second injection (**Supplementary Figure 1**) targeted the PCC sulcus, and also resulted in labeling around the medial orbital sulcus, although it was less specific. We concluded that although the PCC does connect with other OFC subareas (OFC*out*), its relationship with the subareas surrounding the medial orbital sulcus (from here on referred to as OFC*in*) is unique. We next sought to examine the functional properties of this circuit.

**Figure 1.**
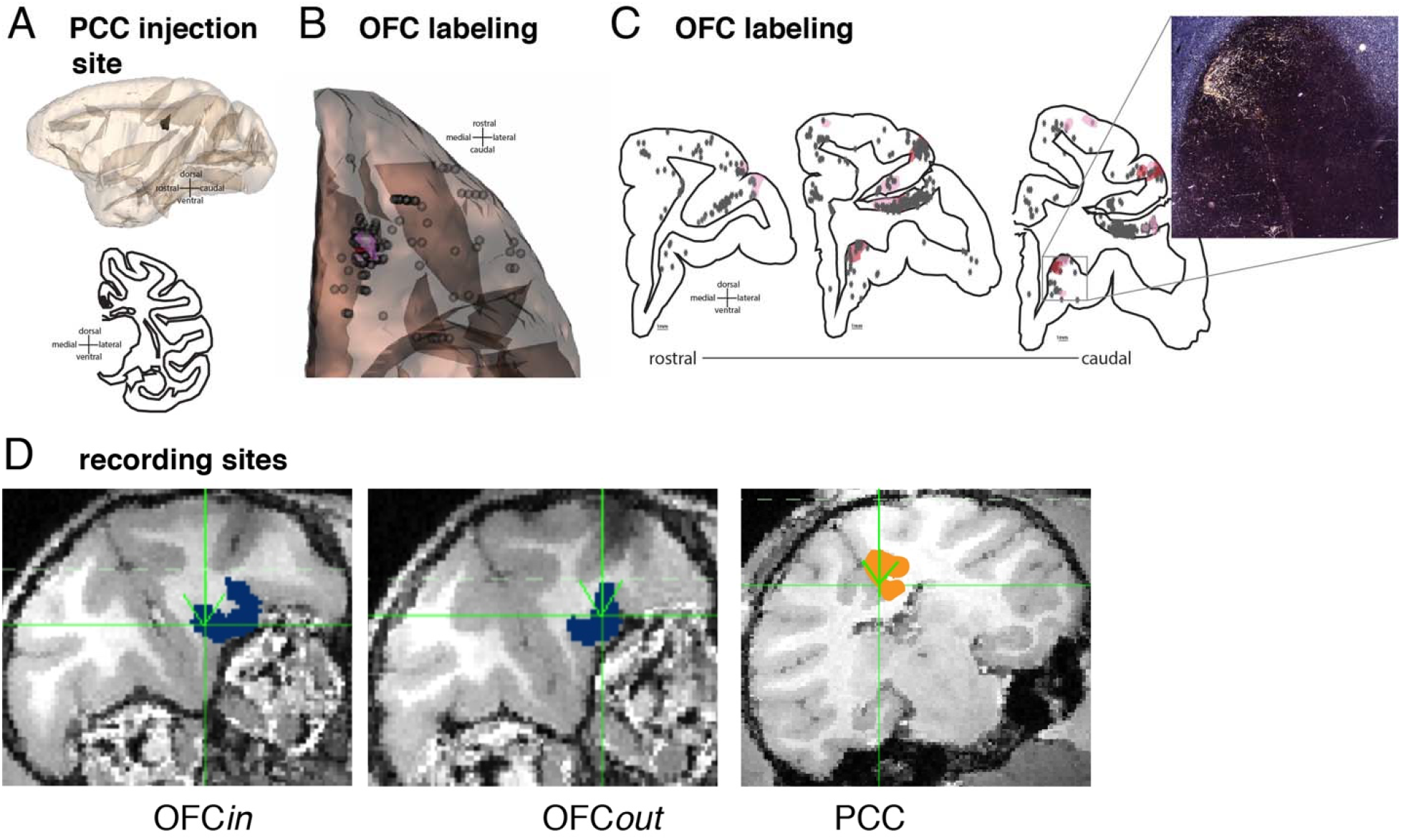
Anatomical connectivity between OFC and PCC and matching recording sites. **A.** Injection site (black, Case M1FR) is rendered in 3D and shown in a sagittal view (top) and on a coronal slice (bottom). **B.** Projections to the OFC rendered in 3D and shown on an orbital view. Red indicates dense terminal fields; pink indicates light terminal fields; gray spheres are labeled cells. The majority of OFC labeling is around the medial orbital sulcus. **C.** Coronal slices with full PFC labeling, colors are as in (B). A photomicrograph indicates label around the medial orbital sulcus. **D**. Coronal sections of example recording site from each of OFC (dark blue colored region), with OFC*in* on left and OFC*out* in the middle, and PCC (orange colored region).

### Behavior and electrophysiology

We recorded neural activity in all three regions--PCC, OFC*in*, and OFC*out* (**Figure 1D**)--while rhesus macaques (*Macaca mulatta*, Subjects P and S) performed a well-established economic choice task (Strait et al., 2014; Farashahi et al., 2018; **Figure 2A**). The critical features of the task are its asynchronous presentation of options (offer 1 and offer 2) and the random order of presentation of options by location (left vs. right), which allowed us to examine the relationship between encodings of offer both abstractly (by time of presentation) and concretely (by side of presentation). As in our past studies using this task (e.g. Strait et al., 2014), both subjects reliably chose the option with higher expected value, indicating high choice accuracy (**Supplementary Figure 2**). We recorded neural ensembles with multiple linear probes simultaneously in both PCC (n=213 neurons) and OFC (n=98 neurons, 44 in OFC*in* and 54 in OFC*out*).

**Figure 2.**
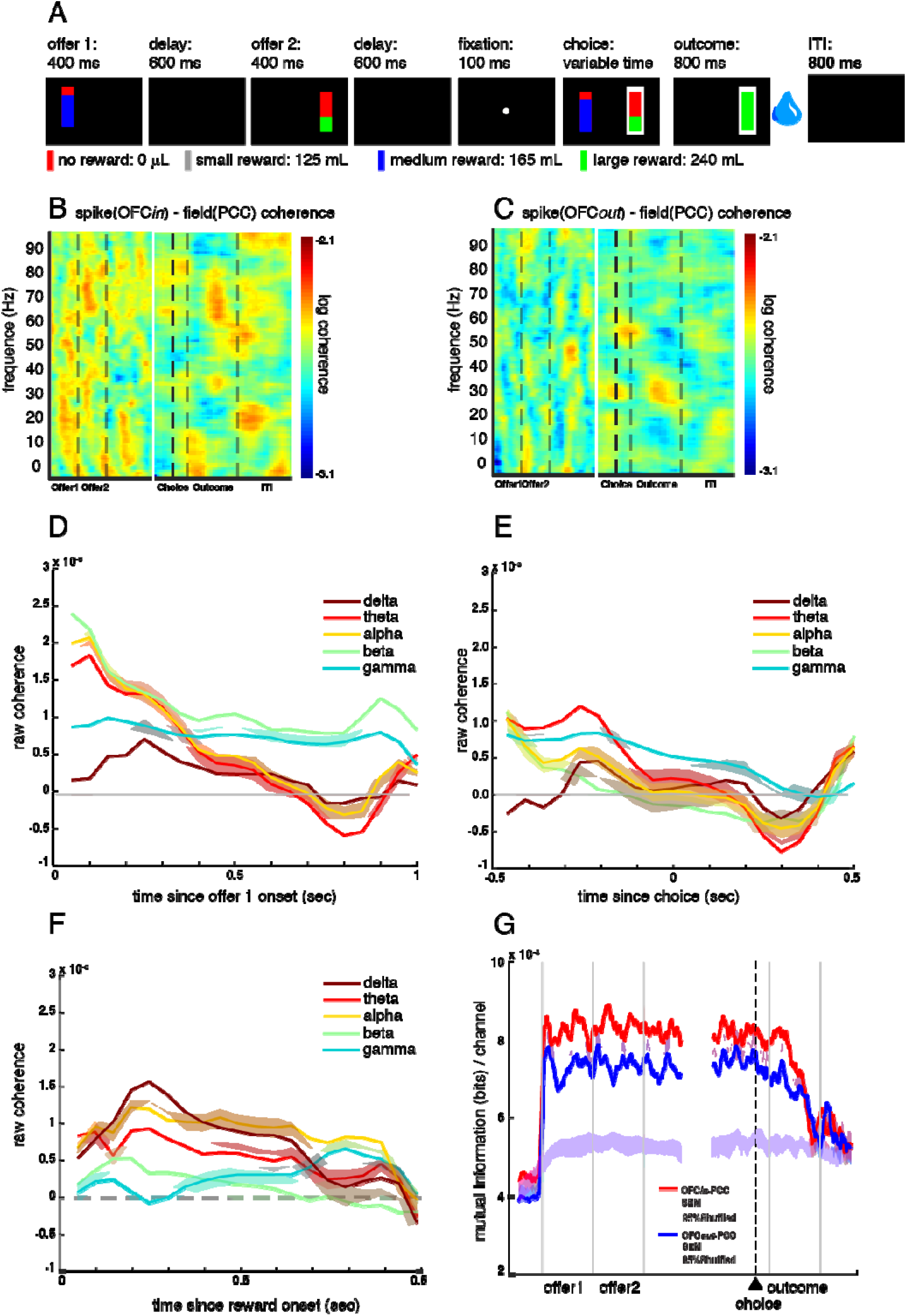
Task and functional connectivity. **A.** Two-option risky choice task. Black rectangles symbolize various task epochs subjects experience during task. Stakes are represented as different colors: small (gray), medium (blue), or large (green) reward. Losing the gamble (no reward) is represented in red. The height of the stakes-color region represents the probability of winning the gamble, and the height of the red-color region represents the probability of losing the gamble. The white frame around the right option in the choice epoch represents the scenario where the subject chooses the right option with eye fixation. The water droplet symbol indicates that reward delivery (or lack thereof) occurs. **B.** Trial-averaged spike-field coherence in OFC*in*_spk_-PCC_lfp_ circuit. X axis: time in a trial. Y axis: frequency. Color: strength of spike-field coherence on log10 scale (warmer colors=higher coherence). Data from the first half of the trial (offer period) was aligned at offer 1 onset. Data from the second half of the trial (choice period) was aligned at choice execution. **C.** Spike-field coherence in OFC*out*_spk_-PCC_lfp_. Conventions as in (B). **D-F**. Difference in spike-field coherence between the two circuits (coherence in OFC*in*_spk_-PCC_lfp_ circuit minus coherence in OFC*out*_spk_-PCC_lfp_ circuit), broken down into different frequency bands as a function of time (**Methods**), during (C) offer 1 epoch, (D) choice epoch, and (E) reward epoch. **G**. Mutual information (averaged across number of channels) in OFC*in*-PCC and OFC*ou*t-PCC circuits. SEM: standard error of the mean. Red shaded area: SEM of mutual information in OFC*in*-PCC circuit. Blue shaded area: SEM of mutual information in OFC*ou*t-PCC circuit. Magenta and cyan shaded areas: the middle 95% range of the randomly shuffled mutual information (500 times) for OFC*in*-PCC and OFC*ou*t-PCC circuits, respectively. Thus, the original (non-shuffled) mutual information values outside of the shaded area is significantly higher/lower than expected by chance.

### Functional connectivity

We asked whether the OFC*in*-PCC circuit shows greater *functional* (rather than anatomical) connectivity than the OFC*out*-PCC circuit. We employed spike-field coherence, which relates the recorded action potentials of one region to the local field potential (LFP) oscillations of another (Buzsáki, 2004; Dal Monte et al., 2020; Buzsaki & Draguhn, 2004; Pesaran, 2010; Scherberger et al., 2005; Widge et al., 2019; see **Methods** and **Supplementary Figure 3**). We found that broadband spike-field coherence between OFC*in* (spikes) and PCC (LFPs) is stronger than coherence between OFC*out* and PCC. Specifically, during the offer epoch, the broadband spike-field coherence in the OFC*in*_spk_-PCC_lfp_ circuit is higher than that in the OFC*out*_spk_-PCC_lfp_ circuit (z=5.01, p<0.001, Wilcoxon signed rank test, **Figure 2B-C**). This effect appears to be broadband; it is significant within all five bands that we tested: delta, theta, alpha, beta, and gamma (**Figure 2D)**. The same pattern occurs in the choice and outcome epochs (OFCin_spk_-PCC_lfp_ > OFC*out*_spk_-PCC_lfp_; choice: z=2.81, p=0.005; outcome: z=3.70, p=0.005). During choice, higher coherence occurs within the theta, alpha, and gamma bands, but not the delta or beta bands (**Figure 2E**). During outcome, higher coherence occurs in all but the beta band (**Figure 2F**).

**Figure 3.**
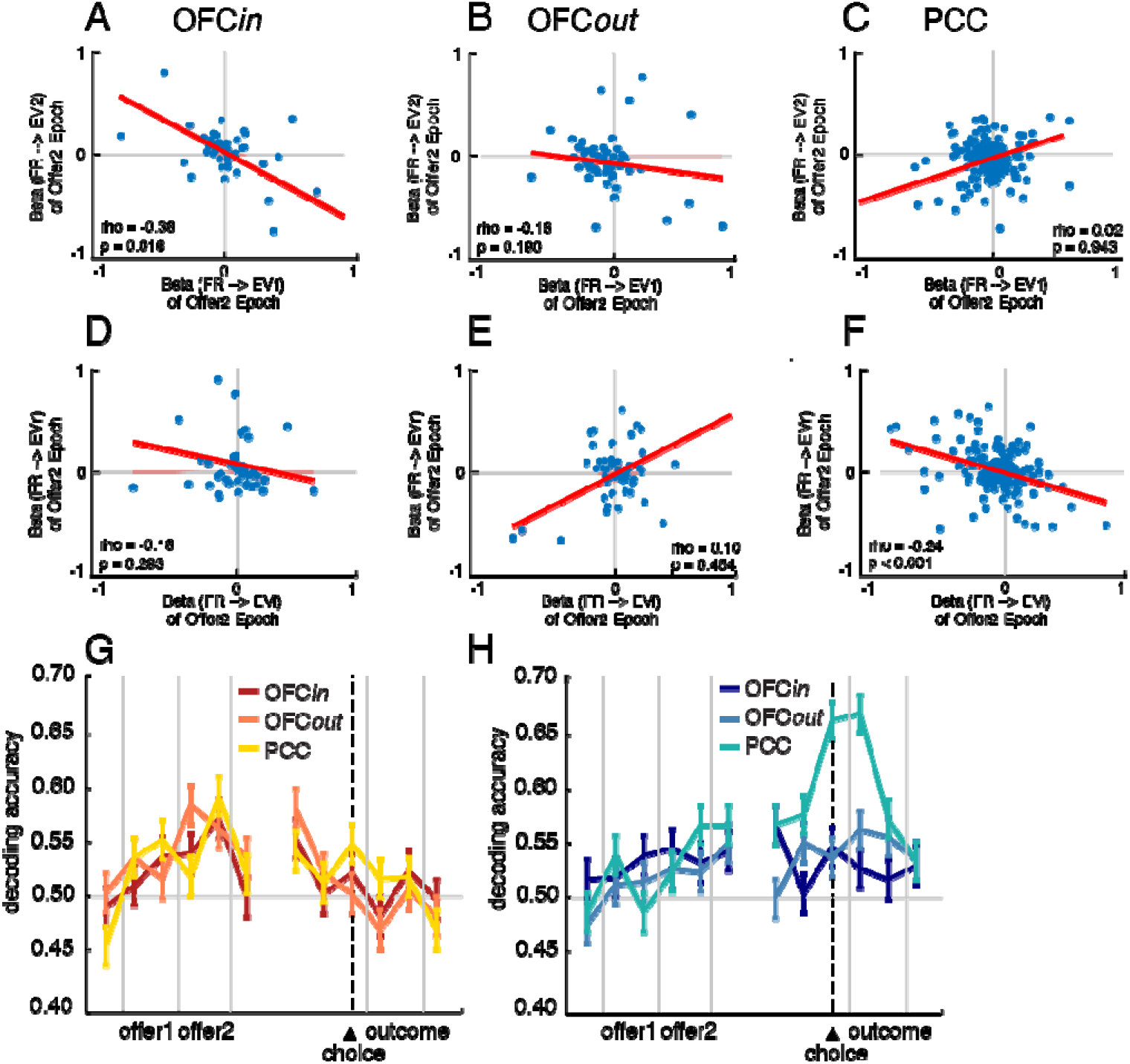
Neural computations. **A-F**: Scatter plots demonstrating population spreads for regression coefficients. Each dot represents one neuron; abscissa and ordinate represent regression coefficients for distinct (and uncorrelated) regressions. Shaded area: 95% confidence interval. **A-C**: Putative mutual inhibition effects (Strait et al., 2014). Y-axis indicates regression coefficient for expected value of offer 2 regressed against firing rate in epoch 2. X-axis indicates regression coefficient for expected value of offer 1 against firing rate in epoch 2. **D-F**: Putative mutual inhibition effects for space (new analysis developed for this project): Y-axis: regression coefficient for expected value of right offer against firing rate in epoch 2. X-axis: regression coefficient for expected value of left offer against firing rate in epoch 2. **A,D:** OFC*in*. **B,E**: OFC*out*. **C,F**: PCC. **G-H**: Decoding accuracy of choice (G is accuracy for offer 1 vs offer 2; H is accuracy for left vs right) based on firing rates using linear discriminant analysis. Y-axis: probability of decoding correctly. X-axis: time in a trial. Error bar: standard error of the mean.

**Figure 4:**
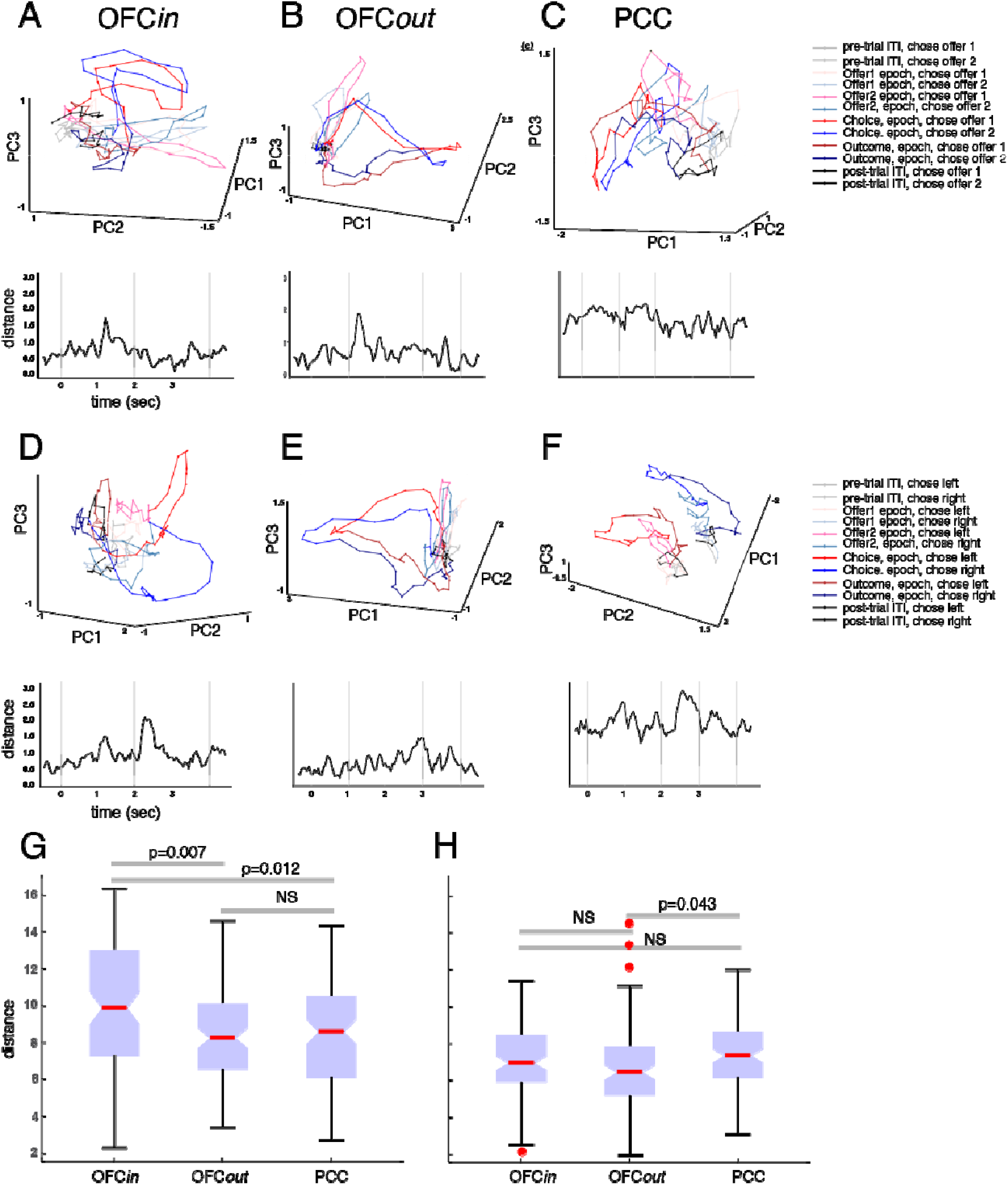
Top plots: trial averaged population activity projected onto top-N PC space (only top-3 PCs are shown here), separated by choice option (offer 1 vs 2) (**A-C**) or choice location (**D-F**), in OFC*in* (left column), OFC*out* (middle column), and PCC (right column). Warm colors: trial averaged population activity for choosing offer 1 (**A-C**) or left offer (**D-F**). Cool colors: trial averaged population activity for choosing offer 2 (a-c) or right offer (d-f). Colors indicate each of the following epochs: the ITI before the current trial, the offer 1 epoch, offer 2 epoch, choice epoch, outcome epoch, and the ITI after the current trial. Bottom plots: separation measured by Euclidean distance between averaged population trajectories (warm and cool colored lines). Y-axis: Euclidean distance. X-axis: time in a trial. Dark line: distance between trial-averaged trajectories for choosing offer 1 vs. offer 2 (**A-C**) or choosing left vs. right offer (**D-F**). Shaded area: middle 95% trial-averaged Euclidean distance between population trajectories from condition-shuffled data. Shuffle was only based on the choice of offer 1 or offer 2 (**A-C**) or on the choice of left or right offer (**D-F**), the cell identities and temporal orders were not shuffled.

We next probed information exchange within our two newly identified circuits by comparing mutual information within each one (see **Methods**). We computed channels as the set of all possible pairs of trains from across the two regions (Timme & Lapish, 2018). Thus, we identified 9372 channels in the OFC*in*-PCC circuit and 11502 channels in the OFC*out*-PCC circuit and calculated the averaged mutual information per channel within each circuit. We found that the OFC*in*-PCC circuit shares higher mutual information than OFC*out*-PCC (7.44×10^−4^ vs. 6.72×10^−4^ bits/channel; z=17.47, p<0.001, Wilcoxon signed rank test). Mutual information in both circuits increased significantly at task onset (p<0.025, shuffle test, see **Supplement**), suggesting that the observed mutual information effect reflects task-driven, rather than spontaneous, fluctuations (**Figure 2G**).

### Neural computation

Functional connectivity results do not speak to the *content* of the information transmitted. We therefore analyzed encoding of task variables with a multiple linear regression model. All three regions encoded offer and outcome values in their respective epochs with similar proportions of neurons, encoding strengths, and latency (**Supplement**). They also all encoded the chosen option (offer 1 vs. 2) and chosen location (left vs. right). However, OFC*in* encoded the chosen option (offer 1 vs 2) with shorter latency (90 ms, F=3.35, p=0.037, GLM Gamma distribution; **Methods**) than both OFC*out* (170 ms, t=−2.14, p=0.033) and PCC (150 ms, t=−2.36, p=0.019), suggesting chosen option information arises first in OFC*in.* PCC appears to be more spatially sensitive than either OFC region: it showed a higher proportion of neurons encoding chosen location than chosen option (χ^2^=5.31, p=0.021, chi-square test); neither OFC region shows this pattern (**Supplement**). PCC and OFC*in* also encoded the chosen location with significantly shorter latencies than OFC*out* (F=5.71, p=0.004; **Supplement**), suggesting that PCC and OFC*in*, but not OFC*out*, negotiate chosen location encoding.

We next examined the negative correlation of regression coefficients for the two offers when offer 2 was revealed, a putative neural signature of value comparison (Azab & Hayden, 2017; Strait et al., 2014). We performed this analysis using a 200-ms analysis window (350 ms after offer 2 onset; the same window identified by the Granger analysis, see below) and found that OFC*in* showed this putative mutual inhibition signal (r=−0.36, p=0.016, Spearman correlation; **Figure 3A**). We did not observe such an effect in OFC*out* (r=−0.18, p=0.190; **Figure 3B**) or in PCC (r=0.02, p=0.943; **Figure 3C**). We also did not find this negative correlation during the later choice epoch (from 400 ms to 200 ms before choice action) in any of the three regions (**Supplementary Figure 4A-C**). The effect size of these negative correlations was not significantly different in OFC*in* vs. OFC*out* (z=−0.93, p=0.176; Fisher’s Transformation test) but was significantly larger in OFC*in* than in PCC (z=−2.32, p=0.010). These results suggest that both OFC subregions, moreso than PCC, were involved in value comparison between offer 1 vs. offer 2 in a presentation order frame, although the effect did not reach significance in OFC*out* alone.

To gain insight into the question of how effector-independent (order-based) value signals are transformed into effector-dependent (spatial-based) ones, we next asked whether regression coefficients for *left* and *right* offer values (EV*l* and EV*r*; as opposed to first and second as in the previous analysis) were negatively correlated. In other words, relaying the mutual inhibition signal to this framework would indicate that neurons carry a decision variable that could potentially be read out by downstream motor areas to guide actions. We found this negative correlation between EV*l* and EV*r* during the offer 2 epoch in PCC (r=−0.24, p<0.001; **Figure 3F**), but not significantly in OFC*in* (r=−0.16, p=0.293; **Figure 3D**) or OFC*out* (r=0.10, p=0.454, **Figure 3E**). Interestingly, the effect size of these negative correlations was not significantly different in OFC*in* vs. PCC (z=−0.49, p=0.313) but was significantly larger in PCC than in OFC*out* (z=−2.21, p=0.014). During the later choice epoch, we found the same signal in both PCC (r=−0.19, p=0.006) and OFC*in* (r=−0.33, p=0.029), but not OFC*out* (r=0.31, p=0.022) (**Supplementary Figure 4D-F**). These results suggest that PCC and OFC*in*, but not OFC*out*, were involved in value comparison between left vs. right offers.

To help understand whether the OFC*in*-PCC circuit transforms the mutual inhibition signals from a value-based comparison to an action-based comparison, we measured Granger causality between time series of mutual signals in OFC*in* and PCC calculated with a 200-ms sliding window (see **Methods**). We found that the strength of mutual inhibition for EV1-EV2 in OFC*in* Granger-caused the strength of mutual inhibition for EV*l*-EV*r* in PCC (gc=40.56, p=0.019), with a 240 ms (4.17 Hz) lag. In the reverse direction, the strength of mutual inhibition for EV*l*-EV*r* in PCC Granger-caused the strength of mutual inhibition for EV1-EV2 in OFC*in* (gc=59.75, p=0.014), but with a much longer lag (380 ms; 2.63 Hz). In contrast, the strength of mutual inhibition for EV1-EV2 in OFC*out* did not Granger-cause the strength of mutual inhibition for EV*l*-EV*r* in PCC with any time lag (see **Methods** for controls for confounding variables). These results suggest that through the communication in the OFC*in*-PCC circuit, but not the OFC*out*-PCC circuit, the computation for value comparison transformed from value space (in OFC*in*) to action space (in PCC).

If the previous result holds, then we would expect to decode choice signal more strongly in value space (in the format of chosen option, offer 1 vs. 2) in OFC*in* but decode choice more strongly in action space (in the format of chosen location, L vs. R) in PCC. We tested for this using Linear Discriminant Analysis (LDA). Although chosen options (offer 1 vs. 2), chosen location (left vs. right), and EV1 (high vs. low) were all significantly decodable from all three regions, PCC indeed showed a significantly higher decodability for chosen location (χ^2^=8.12, p=0.004; **Figure 3G-H; Supplementary Figure 5**). More importantly, we found that the decodability for chosen option (offer 1 vs. 2) in OFC*in* Granger-caused the decodability for chosen location (left vs. right) in PCC (gc=11.19, p=0.025) with a 200 ms (5 Hz) lag. This Granger-causal relation was absent on error trials (gc=3.04, p=0.552). In the reverse direction, the decodability for chosen location (left vs. right) in PCC Granger-caused the decodability for chosen option (offer 1 vs. 2) in OFC*in* (gc=17.59, p=0.025), but with a longer lag (400 ms; 2.5 Hz). In contrast, the decodability for chosen option (offer 1 vs. 2) in OFC*out* did not Granger-cause the decodability for chosen location (left vs. right) in PCC at any time lag (see **Methods** for controls for confounding variables). These results suggest that the OFC*in*-PCC circuit, but not the OFC*out*-PCC circuit, mediates the transformation of choice readout from a value-based to an action-based framework. Speculatively, this transformation may be important for correct choice behavior, since the both the decodability for choice and the Granger causal relation between OFC*in* and PCC was disrupted in error trials (**Supplement**).

We asked whether the population activity dynamics (Afshar et al., 2011; Bartolo & Averbeck, 2020; Churchland et al., 2012; Mante et al., 2013; Yoo and Hayden, 2020) also reflect the translation of choice from value space to action space in the OFC*in*-PCC circuit. Research in motor generation has found that population activity dynamics in the premotor area during the preparatory period determined the possible range of neural dynamics in the primary motor area (M1), and this range determined what hand motion can be generated in M1 (Afshar et al., 2011). To test whether this dynamical generative process of local neural computation could help explain the relayed choice dynamics from abstract value space in OFC*in* to concrete action space in PCC, we conducted PCA on trial-averaged population states for each region and then projected the trial-averaged population activity onto the top-N principal component (PC) space that cumulatively explained > 70% of the variance (**Methods**; we developed this approach in Wang & Hayden, 2017). The projected population trajectories reflect the generative temporal evolution of population dynamics (**Figure 4A-F**), and the separation between trajectories, which distinguished task parameters, became significantly higher than shuffled chance level as the trial unfolded (bottom shaded area). These distinctions diminished in error trials (**Supplementary Figure 6**), suggesting that the population dynamics and their separation are indeed crucial for generating correct choice behavior.

We then projected the trial-by-trial population states onto this top-N PC space to obtain trial-by-trial population trajectories and used adjusted distance to measure the trajectory separation (**Methods**; Murray et al., 2017). We found significantly larger trajectory separation for chosen option (offer 1 vs. 2) in OFC*in* (χ^2^=11.51, p=0.003, Kruskal-Wallis test with Tukey-Kramer multiple comparison) than in OFC*out* (OFCin>OFCout: p=0.007) and PCC (OFCin>PCC: p=0.012; no significant difference between OFC*out* and PCC, p=0.988). This result highlights the specific role of OFC*in* in mediating abstract comparison.

In contrast, we found significantly larger trajectory separation for chosen location (left vs right) in PCC (χ^2^=6.27, p=0.043, Kruskal-Wallis test with Tukey-Kramer multiple comparison) than in OFC*out* (PCC>OFC*out*: p=0.043) but not in OFC*in* (PCC≈OFC*in*: p=0.829; there was no significant difference between OFC*in* and OFC*out,* p=0.164). There was also no such cross-region distinction for EV1 (high vs. low; Supplement; **Supplementary Figure 6**). The trajectory separation differences for chosen option and chosen location were also absent in error trials (**Supplement**), consistent with the intuitive idea that the areal difference in the unfolding trajectory separation contributes to correct choice behavior.

The separation between population trajectories for chosen option (offer 1 vs 2) in OFC*in* Granger-caused the separation between population trajectories for chosen location (left vs. right) in PCC (gc=9.98, p=0.019), with a 150 ms (6.67 Hz) lag. In the reverse direction, the distance between population trajectories for chosen location (left vs. right) in PCC Granger-caused the distance between population trajectories for chosen option (offer 1 vs. 2) in OFC*in* (gc=17.28, p=0.016) but with a much longer lag (350 ms; 2.86 Hz). Interestingly, this “feedback” influence seems to amplify the OFC*in* to PCC input 300 ms after the first instance of Granger causal influence, by increasing the Granger-causality from OFC*in* to PCC (gc=38.29, p<0.001; lag=450ms; 2.22 Hz). In contrast, the distance between population trajectories for chosen option (offer 1 vs. 2) in OFC*out* did not Granger-cause the distance between population trajectories for chosen location (left vs. right) in PCC with any time lag (see **Methods** for the control for confounding variables). These results suggest that while local neural computation was generating choice representations, their unfolding population dynamics also interact with the generative dynamics in other regions. We found the dynamics in OFC*in*-PCC, but not those in OFC*out*-PCC, morphed from developing the separation for choice in value space to developing the separation for choice in action space.

Euclidean distance (i.e. separation; dark line) beyond the shaded area is significant (p<0.05). Specifically, the distance (dark line) larger than (above) the shaded area is where separation between population trajectories is significantly larger than expected by chance (p<0.025). These significant portions mark when the population activity dynamics significantly reflected the choice of offer 1 or offer 2 (**A-C**) or on the choice of left or right offer (**D-F**). **G-H**: Bottom: ranked trial-by-trial adjusted distance. Kruskal-Wallis box plot. The red horizontal line: the median. The bottom and top edges of the box: the 25th and 75th percentiles. The whiskers extend to the most extreme data points not considered outliers. ‘+’ individual outliers.

## DISCUSSION

Here we report the existence of two functionally distinct subregions within the OFC that can be differentiated by their connectivity with the PCC, both anatomically and functionally. OFC*in,* located on the banks of the medial orbital sulcus, seems to have stronger anatomical connectivity with PCC than OFC*out*, which for our electrophysiological recordings was situated lateral to OFC*in.* This boundary seems to correspond to a functional separation that relates to the negotiation between abstract (goods-based) and motor (action-based) modalities. The abstraction transformation is mediated by an OFC*in*-PCC circuit (that is anatomically and functionally connected) and is uncoverable using analyses of value comparison, decoding, and population dynamics. Crucially, instead of copying the more abstract choice signal from OFC*in*, computation within PCC (perhaps with assistance from other input structures) adopts a spatial framework, which allows it to compare and represent the choice in a more concrete, action-based manner. This influence is also bidirectional, with a later PCC (possibly feedback) influence that relays choice from action space back to value space in OFC*in,* and OFC*in* in turn exerts an even stronger influence of relaying choice from value space again to action space in PCC.

We speculate that OFC*in*-PCC forms a bidirectional, mutually excitatory circuit. Our data support the hypothesis that within both regions, a mutually inhibitory local circuit exists to compare offers - in value space in OFC*in* but in action space in PCC. This circuit potentially locks its computation with theta and delta band oscillations to translate choice representation from abstract value space to concrete action space. Moreover, we did not see the information relay between OFC*in* and PCC in error trials, suggesting that the transformation of choice in the OFC*in*-PCC circuit is essential for generating a correct choice. Presumably, after the relay of information between OFC*in* and PCC, a downstream area could use the action-bounded choice signal to form an action plan.

Searching for reward signals in the brain leads to an embarrassment of riches (Vickery et al., 2011; Rushworth et al, 2011; O’Doherty, 2014). The abundance of value is itself mysterious – why would the brain have so many seemingly redundant signals? One possibility is that different value correlates have subtly different roles. That is, they may help negotiate the transform from abstract to concrete spaces in different ways. Our results point to one possible case of this distinction, where some OFC value signals are relatively abstract and others are relatively concrete, but the concrete (motoric) aspects of OFC signals are derived from more specialized PCC signals. More speculatively, our results suggest that even apparent intra-areal redundancy of function may mask an underlying heterogeneity of function.

## METHODS

### Neuroanatomy studies

We injected the bidirectional tracer fluororuby into the PCC of two adult male rhesus macaque (*Macaca mulatta*) subjects. In one (M1FR), the injection site was located at the border of areas 23 and 30 (with some involvement of area 29). In another (M6FR), the injection site was located at the border of areas 23 and 31. We note that, although the PCC is often defined as areas 23 and 31, with areas 29 and 30 instead defined as retrosplenial cortex (Vogt et al., 2006; Leech et al., 2011), we were interested in the functionality of this entire caudal cingulate region. Thus, like some prior authors (Armstrong et al., 1986; Zilles et al., 1986; Mitelman et al., 2005; Vogt et al., 1992), here we defined PCC as areas 23, 31, 29, and 30.

Prior to surgery, anatomical T1 and T2-weighted MRIs (3T for M1FR and 10.5T for M6FR) were obtained at University of Minnesota’s Center for Magnetic Resonance Research. Stereotaxic earbars were filled with Vitamin E solution to visualize on the MRI and guide tracer placement relative to stereotaxic zero.

On the day of surgery, monkeys were tranquilized by intramusculuar injections of ketamine (10mg/kg), midazolam (0.25mg/kg) and atropine (0.04mg/kg). A surgical plane of anesthesia was then maintained via the administration of inhalation of isofluorane (1-3%). Monkeys were placed in a stereotaxic instrument (Kopf Instruments), a midline scalp incision was made, and the muscle and fascia were displaced laterally to expose the skull. A craniotomy (~2-3cm^2^) was made over the PCC, and small dural incisions were made only at injection sites. Both monkeys received injections of FR (50nl, 10% in 0.1M PB, pH 7.4, Invitrogen) in the PCC, as well as injections of additional tracers (lucifer yellow, fluorescein, wheat germ agglutinin conjugated to horseradish peroxidase) in other regions not described here. These do not cross-react with FR and were made distant from the PCC site. Tracers were pressure-injected over 10 min using a 0.5-μl Hamilton syringe. Following each injection, the syringe remained in situ for 20–30 min. Twelve to 14 days after surgery, monkeys were again deeply anesthetized and perfused with 4L of saline followed by 6L of a 4% paraformaldehyde/1.5% sucrose solution in 0.1 M PB, pH 7.4. Brains were postfixed overnight and cryoprotected in increasing gradients of sucrose (10, 20, and 30%). Serial sections of 50 μm were cut on a freezing microtome into cryoprotectant solution.

One in eight sections was processed free-floating for immunocytochemistry to visualize the tracer. Tissue was incubated in primary anti-FR (1:6000; Invitrogen) in 10% NGS and 0.3% Triton X-100 (Sigma-Aldrich) in PO4 for 4 nights at 4°C. After extensive rinsing, the tissue was incubated in biotinylated secondary antibody followed by incubation with the avidin-biotin complex solution (Vectastain ABC kit, Vector Laboratories). Immunoreactivity was visualized using standard DAB procedures. Staining was intensified by incubating the tissue for 5–15 s in a solution of 0.05% DAB tetrahydrochloride, 0.025% cobalt chloride, 0.02% nickel ammonium sulfate, and 0.01% H2O2. Sections were mounted onto gel-coated slides, dehydrated, defatted in xylene, and coverslipped with Permount.

Using a Zeiss M2 AxioImager, light microscopy was used to outline brain sections, PCC injection sites, frontal cortical terminal fields, and frontal cortical labeled cells on 1 in 24 sections (1.2mm apart). Terminal fields were outlined in darkfield using a 2.0, 4.0, or 10× objective with Neurolucida software (MicroBrightField Bioscience). Terminal fields were considered dense when they could be visualized at a low objective (2.6×) (Haber et al. 2006); otherwise, terminal fields were considered sparse. Thin, labeled fibers containing boutons were marked as terminating; thick fibers without boutons were considered passing. Retrogradely labeled cells were identified under brightfield microscopy (20×) using StereoInvestigator software (MicoBrightField Bioscience).

Cases were registered and rendered in 3D in the following way. For each case, a stack of 2D coronal sections was created from its Neurolucida chartings. This stack was imported into IMOD (Boulder Laboratory for 3D Electron Microscopy, Kremer et al. 1996), and a 3D reconstruction that contained the injection sites, terminal fields, and cells was created for each case separately. To render these and merge cases together, we used a reference model from the NIMH Macaque Template (Seidlitz et al., 2017), imported into IMOD. Placement of all contours—injection sites, terminal fields, cells, area outlines—were assessed according to cortical and subcortical landmarks in the brain, then checked with the original case and corrected as needed.

### Neurophysiology studies

#### Subjects

Two male rhesus macaques (*Macaca mulatta*) served as subjects to the neurophysiology experiment. All animal procedures were approved by the University Committee on Animal Resources at the University of Rochester (neurophysiology studies) and by the Institutional Animal Care and Use Committee at the University of Minnesota (neurophysiology and neuroanatomy studies). The experiments were designed and conducted in compliance with the Public Health Service’s Guide for the Care and Use of Animals. These subjects were used in past studies involving set shifting and risky choice (Sleezer et al., 2016; Pirrone et al., 2018; Heilbronner and Hayden, 2016).

#### Recording Sites

Two Cilux recording chambers (Crist Instruments) were placed over central OFC and PCC (Paxinos et al., 2009; see also Öngür & Price, 2000; Leech & Sharp, 2014; Mufson and Pandya, 1984; Figure 1D). Note that this posterior region is overlapping with but ventral to a region we have previously recorded in known as CGp (Heilbronner et al., 2013; Hayden et al., 2009; Hayden et al., 2010). Position was verified by magnetic resonance imaging with the aid of a Brainsight system (Rogue Research Inc.) for subject P and Cicerone system (Dr. Matthew D. Johnson at University of Minnesota) for subject S. Neuroimaging was performed at the Rochester Center for Brain Imaging, on a Siemens 3T MAGNETOM Trio Tim using 0.5 mm voxels. We confirmed recording locations by listening for characteristic sounds of white and gray matter during recording, which in all cases matched the loci indicated by the Brainsight system or Cicerone system.

#### Recording techniques

Multicontact electrodes (V-probes, Plexon, Inc) were lowered using the same microdrive system until positioned within the OFC. Following a settling period, all active cells were recorded. Electrodes were lowered using a microdrive (NAN Instruments) until the target region was reached. This lowering depth was predetermined and calculated with the aid of either Brainsight or Cicerone system to make sure the majority of the contacts on the V-probe were in the gray matter of the recording region. Individual action potentials were isolated on a Ripple Grapevine system (Ripple, Inc.). Neurons were selected for study solely on the basis of the quality of isolation; we never pre-selected based on task-related response properties. Cells were sorted offline with Plexon Offline Sorter (Plexon, Inc.) by hand by MZW. No automated sorting was used.

#### Eye Tracking and Reward Delivery

Eye position was sampled at 1,000 Hz by an infrared eye-monitoring camera system (SR Research). Stimuli were controlled by a computer running MATLAB (Mathworks) with Psychtoolbox (Brainard, 1997) and Eyelink (Cornelissen et al., 2002) Toolbox. A standard solenoid valve controlled the duration of fluid reward delivery. For part of the behavioral training, subjects received grape juice or cherry coke instead of water as reward. However, water reward was used during all neural recording sessions. The relationship between solenoid open time and water volume was established and confirmed before, during, and after recording.

#### Behavioral task

Subjects performed a two-option gambling task identical to the one we used in a previous investigation (**Figure 1**, Strait et al., 2014; see Heilbronner, 2017 for context). Two offers were presented on each trial. Each offer was represented by a rectangle 300 pixels tall and 80 pixels wide (11.35° of visual angle tall and 4.08° of visual angle wide). Options offered either a gamble or a safe (100% probability) bet for liquid reward. Gamble offers were defined by both reward size and probability, which were selected with uniform probabilities and independently of one another for each offer and for each trial. Each gamble rectangle had two sections, one red and the other either blue or green. The size of the blue or green portions indicated the probability of winning a medium (165 μL) or large reward (240 μL), respectively (**Figure 1**). These probabilities were drawn from a uniform distribution between 0% and 100%. Safe offers (1 out of every 8 offers) were entirely gray, and selecting one would result in a small reward (125 μL) 100% of the time.

Offers were separated from the central fixation point by 550 pixels (27.53° of visual angle). The sides of the first and second offer (left or right) were randomized each trial. Each offer appeared for 400 ms followed by a 600 ms empty screen. After the offers were presented one at a time, a central fixation point appeared, and the monkey fixated on it for 100 ms. Then both offers appeared simultaneously and the animal indicated its choice by shifting gaze to its preferred offer, maintaining fixation on it for 200 ms. Failure to maintain gaze for 200 ms would return the monkey to a choice state. Thus, subjects were in theory free to change their mind if they did so within 200 ms (although they seldom did). Following a successful 200-ms fixation, the gamble was immediately resolved and a liquid reward was delivered. Trials that took more than 7 sec were considered inattentive and were excluded from analysis (this removed <1% of trials). Outcomes that yielded rewards were accompanied by a white circle in the center of the chosen offer (see Figure 1B). Each trial was followed by an 800-ms inter-trial interval (ITI) with a blank screen.

Probabilities were drawn from uniform distributions with resolution only limited by the size of the screen’s pixels, which let us present hundreds of unique gambles. Offer reward sizes were selected at random and independent of one another with a 43.75% probability of blue (medium reward) gamble, a 43.75% probability of green (large reward) gambles, and 12.5% probability of safe offers. Note that this means two offers with the same reward size could be (and often were) presented in the same trial.

#### Statistical analyses: Behavior

Only trials accompanying the recording sessions were analyzed for the current paper. For choice accuracy, we defined the correct choice as choosing the offer with expected value higher than or equal to that of the alternative offer. Expected value (EV) is the product of stakes multiplied by probability of winning (getting rewarded, in contrast to getting no reward). Probability of choosing offer 1 as a function of value difference (EV1-EV2) is fitted with generalized linear with logistic transform function and binomial distribution. The error bars indicate 95% confidence intervals from the logistic regression model.

#### Statistical analyses: Spectral analyses

Local field potentials (LFP) were collected during recording sessions along with spike data using the Ripple Grapevine system. LFP data from each contact of the Plexon v-probes were used. Raw data was low-pass filtered at 100Hz and notch-filtered at 60 Hz. All filtering and frequency-domain (spectral) analyses were conducted in Matlab with Chronux toolbox (Bokil et al., 2010). Power spectra in all three regions were calculated with all LFP channels. Spike-field coherence was calculated using every combination of each spike train in one area and each channel of LFP in another area. Coherence comparison used non-parametric statistics: Wilcoxon signed rank test and Kruskal-Wallis test, both conducted in Matlab. We used the following bandwidths for analyses: Delta (0.5-5 Hz), Theta (5-10 Hz), Alpha (10-15 Hz), Beta (15-30 Hz), and Gamma (>30 Hz). For coherence comparisons, we calculated the coherence with a frequency-resolved method, such that we re-adjusted the sliding calculation window widths to be four times the max length for each frequency band. We aligned data to either offer 1 or choice to achieve a better temporal resolution of the coherence tests.

#### Statistical analyses: Mutual information

Mathematically, mutual information is defined as I[X;Y]=H[X]-H[X|Y]=I[Y;X], where *I* is the mutual information between random variables *X* and *Y*. It quantifies the information *X* gives upon observing *Y* and is the same as the information *Y* gives upon observing *X*. Equivalently, it captures how much uncertainty about *X* decreases after learning *Y*, and vice versa. We used the Neuroscience Information Theory Matlab toolbox to calculate the mutual information between two spike trains, one from each brain area of interest (Timme & Lapish, 2018).

To test whether the mutual information in OFC*in*-PCC or OFC*out*-PCC during task was higher than expected chance, we shuffled each single-unit’s brain area identity to form shuffled ensembles with the same sizes as the original data. Then we shuffled temporal sequences within ITI and, separately, within active task-time. The temporal shuffling is to test whether the increase in mutual information was above chance level and driven by engaging in the task. We then calculated mutual information based on these shuffled ensembles. We repeated this procedure 500 times and obtained the middle 95% range of the shuffled mutual information as a function of time (**Figure 3F**, shaded magenta and cyan for OFC*in*-PCC and OFC*out*-PCC circuits, respectively). Thus, any value outside the shaded area is significantly higher/lower than expected by chance.

#### Statistical analyses: Encoding

We used a sliding multiple linear regression to characterize the encoding of all task variables (stakes, probabilities, expected values of offer 1 and offer 2, chosen option, chosen location, whether offer 1 was presented on left vs. right, and choice outcome [win or lose * stakes]). To do so, we took the normalized firing rates (FR) of each neuron, averaged across a 200 ms time bin, and then regressed against task parameters. The sliding procedure slid forward with a 10 ms step size. For offer epochs, we used a multiple linear regression model with stakes, probabilities, and expected values (EV) as predictors. Expected value is defined as the product of stake and probability. For the rest of the epochs, we used a multiple linear regression model with stakes, probabilities, EV1, EV2, chosen option (offer 1 vs. 2), chosen location (left vs. right), outcome (received outcome, 0 for lost gamble, reward of the stake’s size for won gamble), and whether offer 1 appeared on the left or right side of the screen. For later tests looking the expected value tuning for left and right offers, we used a multiple linear regression model with stakes, probabilities, left EV (EV*l*), right EV (EV*r*), chosen option (offer 1 vs. 2), chosen location (left vs. right), outcome (actually received outcome, 0 for lost gamble, reward of the stake’s size for won gamble), and whether offer 1 (first appeared offer) appeared on the left or right side of the screen. All predictors were centered and converted to categorical variables when applicable. The response variable, firing rates, were normalized for each neuron across trials to avoid spurious correlation (Blanchard and Hayden, 2014).

Proportion of neurons was calculated based on whether neurons significantly encoded a single parameter of interest. Encoding strength was defined as the t-statistics of each predictor variable from the multiple regression. We used t-statistics since they are not influenced by the actual range of each variable (even though we centered all predictor variables) and are comparable across variables. The comparison of encoding strength across all three regions used the nonparametric Kruskal-Wallis test. Latency was defined as, within the analyzed event window, the time lapsed until the encoding strength of the variable of interest reached the peak for each neuron. Then the peak time for a region was calculated as the median of each neuron’s peak time. Latency calculation was based on all neurons and not only the significantly tuned ones. Whether latencies from all three regions were significantly different from one another was tested with generalized linear model (GLM) with a Gamma distribution, due to the fact that timing data, such as latency or reaction time, are better described by a Gamma distribution than a Gaussian distribution.

For mutual inhibition, we took the regression coefficients from the above described multiple regression models for the offer 2 epoch and the choice epoch respectively. Then we correlated the coefficients for offer 1 vs. 2 or EV*l* vs. EV*r* with a Spearman correlation. Spearman correlation is chosen to avoid spurious correlation caused only by a few outliers. The strength of mutual inhibition signal is the Spearman correlation coefficient.

#### Statistical analyses: Granger causality

Granger causality measures how one time series could predict (Granger-cause) another time series, after controlling for the fact that the later time series’s early sequences also predicts its own later sequences (Granger, 1969). Sometimes, calculation of Granger causality is also conditioned on simultaneously observing other potentially confounding time series (Lutkepohl, 2007). For all Granger causality tests, we first used the Augmented Dickey-Fuller test with the autoregressive model with drift variant (ARD) to determine whether a time series was stationary. Then we used the vector autoregression (VAR) model to determine the best time lag to use through model comparison (Akaike information criterion) with different time lags. Then the Granger causality test was used on stationary time series or with a correction for non-stationary time series. All Granger causality analyses in this paper tested the Granger-causal relation between two key variables but also included the conditional term with all other potentially confounding variables. That is, none of the potentially confounding decodability could explain the effect we saw. All Granger causality tests were carried out in Matlab. Matlab functions used: adftest, varm, estimate, summarize, gctest, the Econometrics Toolbox.

#### Statistical analyses: Decoding

We first organized population activity patterns for the training and testing of the linear discriminant analysis (LDA) decoder. For each trial, we aligned the normalized firing rates of each neuron at the onset of offer 1 presentation and took firing from 500 ms before this onset through 2500 ms after this onset as the offer period (including 500 ms ITI before offer 1, offer 1 epoch, offer 2 epoch, and the first 500 ms of decision-making). We also aligned the normalized firing rates of each neuron at choice execution (when eye-fixation on the chosen offer passed 200 ms and thus signaled commitment to the choice). Then we took the FR from 1500 ms before this onset through 1500 ms after this onset as the choice period (including 1500 ms pre-choice, outcome delivery, and ITI). We then slid through the offer and the choice periods and generated non-overlapping population activity patterns that were 50 ms in width and tiled the entire offer and choice periods.

Then we followed a four-fold cross validation procedure, which involved training different LDA decoders on 75% of the correct trials to differentiate the chosen option (offer 1 vs. 2), the chosen location (left vs. right), and the expected value of offer 1 (EV1 high vs. low) on each trial. Then we tested the decoder on the other 25% of the correct trials. Decoding accuracy in error trials was obtained by using the same trained LDA decoders to decode all error trials (since none of the error trials were used for training). For EV1 high vs. low, we compared EV1 from each trial to the mean EV of all offers. If the EV1 was larger than or equal to the mean, then it was counted as a high EV1, otherwise low.

#### Statistical analyses: Population dynamics

To measure the dynamics in population neural activities, we first organized our spiking data into population states. We defined the population state as the normalized firing rate of each of all simultaneously recorded neurons, averaged over a 200 ms time bin, in each region. Then we slid across all time points in each trial with a 50 ms step size to calculate population states at each sliding step. We calculated these series of population states for two sets of simultaneously recorded ensembles in OFC*in*, OFC*out*, and PCC, one from each subject. We then applied principal component analysis (PCA) to identify a lower-dimensional space to then measure the population dynamics. We first selected and grouped all correct trials based on whether (1) offer 1 or offer 2 was chosen; (2) left or right offer was chosen; and (3) offer 1 was a higher or lower than average value of offers. Then we conducted PCA on the trial averaged population states for each pair of the above-mentioned three pairs of conditions. To make the measures of population dynamics comparable across regions, we defined top-N PC space as the top N principal components that captured at least 70% of the variance. For subject P, N equals 6 in OFC<u>*in*</u>, 5 in OFC*out*, and 15 in PCC. For subject S, N equals 3 in OFC*in*, 5 in OFC*out*, and 3 in PCC. We then projected trial-averaged or trial-by-trial population states from correct or error trials and each pair of conditions onto this top-N PC space. This projection resulted in pairs of population trajectories corresponding to pairs of conditions in the top-N PC space expanding the whole trial length. We then measured the Euclidean distance at each time point in a trial between the pairs of population trajectories. We used a shuffle procedure in which trials were shuffled across conditions. This shuffle procedure was implemented 1000 times to generate 1000 randomized trial-averaged trajectories for each trial condition, and significance cutoff were set at the top and bottom 2.5% of the shuffled results. For trial-by-trial population state projections that resulted in a pair of two sets of population trajectories (that is, each trajectory corresponded to a specific trial condition), we calculated the adjusted Euclidean distance. The adjusted Euclidean distance is the Euclidean distance ***across*** conditions (cross distance) normalized by the Euclidean distance ***within*** conditions. Cross distance was defined as the Euclidean distance from one point on one trajectory in one trial condition to all the trajectories’ corresponding time point in the other trial condition. Self distance/dispersion was defined as the Euclidean distance of one point on one trajectory in one trial condition to all the other trajectories’ corresponding time point in the same trial condition.

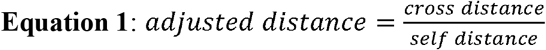

Normalizing the cross distance with self distance controls for the “internal noise level” to make the distance comparable across regions (Murray et al., 2017). The distance, or separation, between population trajectories from pairs of trial conditions represents the population neural activity variance devoted to distinguish those trial conditions (Wang & Hayden, 2017). Intuitively, it can be interpreted as: the larger the distance/separation between trajectories for different conditions, the more information the variance in this neural population conveys to tell these conditions apart. PCA analysis and Euclidean distance calculation used pca and pdist2 functions in Matlab.

## Acknowledgements

We thank Giuliana Loconte, Hannah Lee, Tanya Casta, Mark Grier, Megan Monko, and Adriana Cushnie for experimental help.

## Supplementary material

### Neuroanatomy

A second injection, M6FR, targeted the PCC sulcus. There was anatomical connectivity with OFC*in*, although it was less specific than observed in M1FR (**Supplementary Figure 1**).

**Supplwmentary Figure 1:**
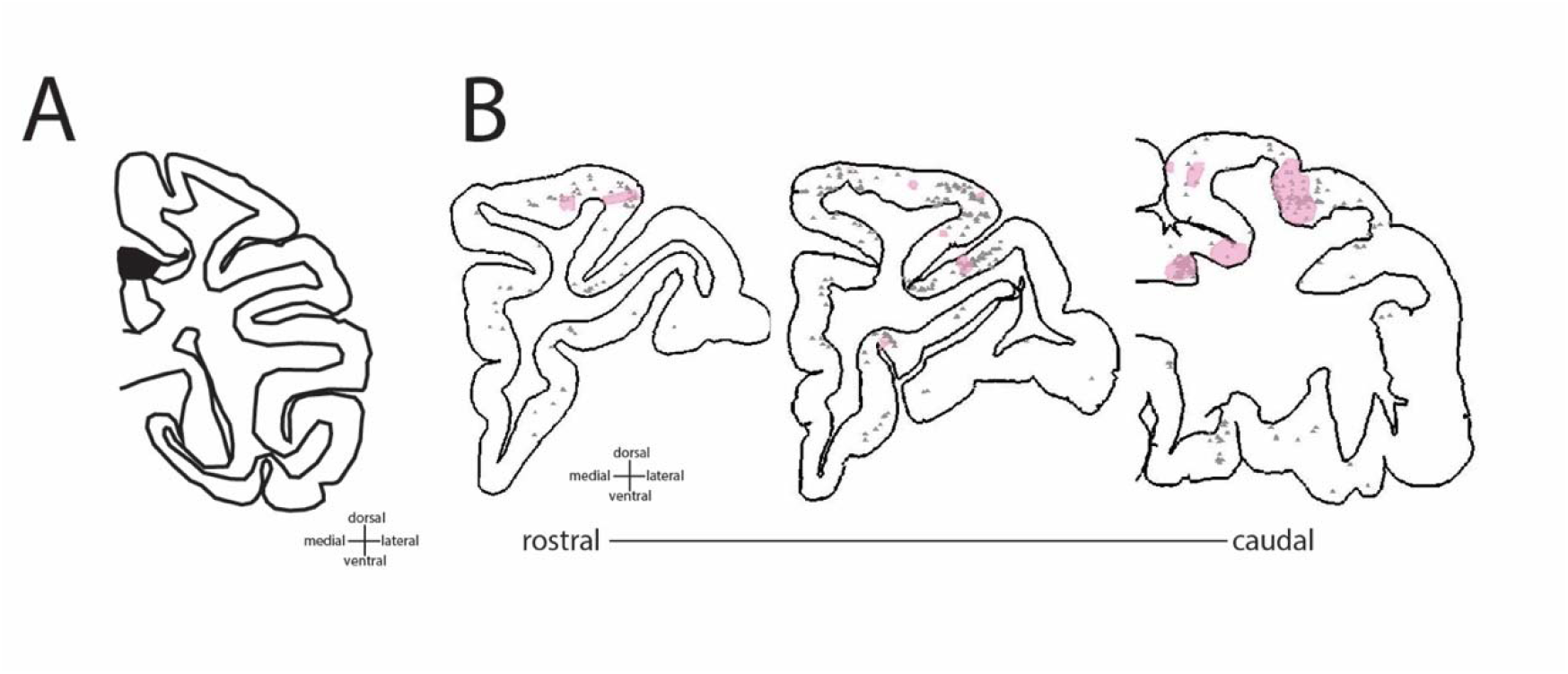
Anatomical connectivity in prefrontal cortex following an injection in the PCC sulcus. **A.** Injection site in PCC **B.** Terminal field labeling shown in pink; retrograde labeling shown as gray dots.

### Behavior of each subject in the gambling task

We examined the behavior of two male macaque subjects (*Macaca mulatta*, subjects P and S) performing a well-studied two-option risky choice task (Strait et al., 2014). The data and results we present here have not been published before, but qualitatively replicate our past findings. Specifically, behavioral data indicate that subjects understood the key elements of the task., They preferred offers with the larger expected value on 73.10% of the trials (for individual subjects, see below). This proportion is significantly higher than expected by chance (p<0.001, binomial test). It is also quantitatively similar to numbers we have found using the same task in other subjects (Strait et al., 2014; Strait et al., 2015). Subjects’ willingness to choose an offer varied as a function of the difference in values between the two offers (**Supplementary Figure 2A-B**). Both subjects slightly preferred offer 2, although the size of the effect was small; choosing offer 1 46.90% of the time). Note that these behavioral results are restricted to trials in which our physiological recordings met criteria for analysis. Data collected in other sessions were not noticeably different (data not shown).

Behaviors of each individual subject closely resembled those of two subjects combined as reported in the main text. Subject P preferred offers with the larger expected value on 73.35% of the trials. This proportion is significantly higher than expected by chance (479 out of 653, p<0.001, binomial test). P shifted choices from offer 1 to offer 2 as the expected value difference of offer 1 minus offer 2 decreased, even with a slight bias against offer 1 (psychometrics function slightly shifted towards right, choosing offer 1 45.18% of the time). Subject S preferred offers with the larger expected value on 72.39% of the trials. This proportion is significantly higher than expected by chance (367 out of 507, p<0.001, binomial test). S shifted their choice from offer 1 to offer 2 as the expected value difference of offer 1 minus offer 2 decreased, even with a slight bias against offer 1 (psychometrics function slightly shifted towards right, choosing offer 1 45.18% of the time).

**Supplwmentary Figure 2:**
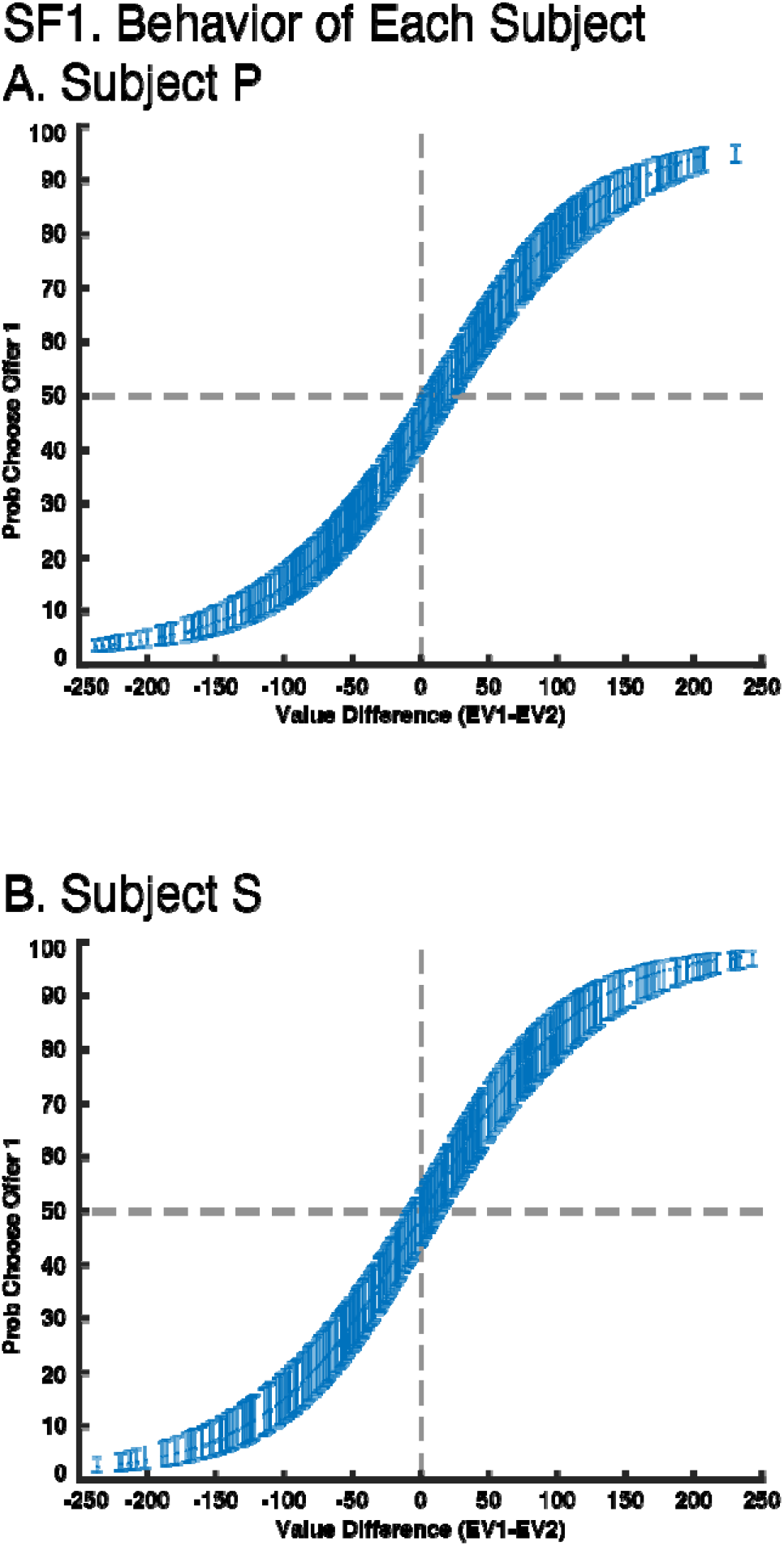
Behavior of Each Subject. **A.** Choices of subject P. **B.** Choices of subject S. EV, expected value (see **Methods**). Gray dotted lines represent visual reference for value 0 on X axis and value 50 on Y axis. Error bars on the fitted sigmoidal function represents 95% confidence interval from the model estimation.

### Functional connectivity

We first characterized the local field potentials in each of the OFC*in*, OFC*out*, and PCC regions. With multitaper spectral analyses, we show that power peaked around 10 Hz in OFC*in* and OFC*out*, and around 10 and 20 Hz in PCC (**Supplementary Figure 3A-C**). It also shows that our notch filter effectively removed power around 60 Hz.

The higher coherence in the OFC*in*_spk_ - PCC_lfp_ circuit was also observed within all specific bands that we tested: the delta (0.5-5 Hz) frequency band (z=2.53, p=0.012), the theta (5-10 Hz) band (z=3.55, p<0.001), the alpha (10-15 Hz) band (z=3.83, p<0.001), the beta (15-30 Hz) band (z=4.38, p<0.001), and the gamma (30-100 Hz) band (z=5.51 p<0.001). Comparing the coherence for each frequency band within each circuit during offer epoch, there was no significant difference among frequency bands within either the OFC*in*_spk_ - PCC_lfp_(χ^2^=3.95, p=0.413, Kruskal-Wallis test) or the OFC*out*_spk_ - PCC_lfp_(χ^2^=2.28, p=0.685, Kruskal-Wallis test) circuit.

We observed similar results in the choice epoch. We also found higher coherence in OFC*in*_spk_ - PCC_lfp_ than in OFC*out*_spk_ - PCC_lfp_ circuit, within the theta (z=1.98, p=0.047), the alpha (z=3.14, p=0.002), and the gamma (z=3.73, p<0.001) bands, although not within the delta (z=1.10, p=0.271) or the beta (z=1.41, p=0.159) bands. Comparing the coherence for each frequency band within each circuit, during choice epoch, there was no significant difference among frequency bands within either OFC*in*_spk_ - PCC_lfp_ (χ^2^=1.81, p=0.771, Kruskal-Wallis test) or OFC*out*_spk_ - PCC_lfp_ (χ^2^=2.53, p=0.640, Kruskal-Wallis test).

Finally, we observed the same general pattern during the outcome epoch. We also found higher coherence OFC*in*_spk_ - PCC_lfp_ than in OFC*out*_spk_ - PCC_lfp_ circuit, within the delta (z=3.36, p<0.001), the theta (z=2.87, p=0.004), the alpha (z=3.70, p=0.002), and the gamma (z=2.05, p=0.040) bands, although not within the beta (z=1.27, p=0.204) band. Comparing the overall coherence for each frequency band during reward epoch, there was significant difference among frequency bands within OFC*in*_spk_ - PCC_lfp_ (χ^2^=14.32, p=0.006, Kruskal-Wallis test with Tukey-Kramer multiple comparison) circuit. Specifically, within OFC*in*_spk_ - PCC_lfp_, the coherence in the beta band was significantly lower than that in the theta band (p=0.021) and that in the alpha band (p=0.009). Similarly, comparing the overall coherence for each frequency band during the reward epoch, there was a significant difference among frequency bands within OFC*out*_spk_ - PCC_lfp_ circuit (χ^2^=17.15, p=0.002, Kruskal-Wallis test with Tukey-Kramer multiple comparison). Specifically, within OFC*out*_spk_ - PCC_lfp_, the coherence in the alpha band was significantly lower than that in the theta band (p=0.007) and that in the gamma band (p=0.029).

Together, we found greater coherence in OFC*in*_spk_ - PCC_lfp_ than in OFC*out*_spk_ - PCC_lfp_, suggesting stronger functional connectivity. This pattern of enhanced coherence was not found in the reverse direction (that is, in the PCC_spk_ - OFC*in*_lfp_, PCC_spk_ - OFC*out*_lfp_ circuits) or their comparison (**Supplementary Figure 3D-J**).

We further compared the broadband spike-field coherence in the reverse direction to that reported in the main text. We found significantly higher broadband coherence in OFC*in*_spk_ - PCC_lfp_ than PCC_spk_ - OFC*in*_lfp_ (z=4.83, p<0.001, Wilcoxon signed rank test, **Supplementary Figure 3D**). The broadband coherence was also higher in OFC*out*_spk_ - PCC_lfp_ than in PCC_spk_ - OFC*out*_lfp_ (z=2.90, p=0.004, Wilcoxon signed rank test, **Supplementary Figure 3E**). We also found significantly higher broadband coherence in PCC_spk_ - OFC*in*_lfp_ than in PCC_spk_ - OFC*out*_lfp_ (z=2.76, p=0.006, Wilcoxon signed rank test; **Supplementary Figure 3D-E**) but no significant differences in broadband coherence between OFC*in*_spk_ - OFC*out*_lfp_ and OFC*out*_spk_ - OFC*in*_lfp_ (z=0.15, p=0.883, Wilcoxon signed rank test; **Supplementary Figure 3F-G**).

Spike-field coherence is theorized to capture long-range input from the spiking region to the field region. Our results suggest that the enhanced synchronization for OFC*in*-PCC could be dominated by OFC*in*’s input to influencing PCC local neurocomputation.

**Supplementary Figure 3:**
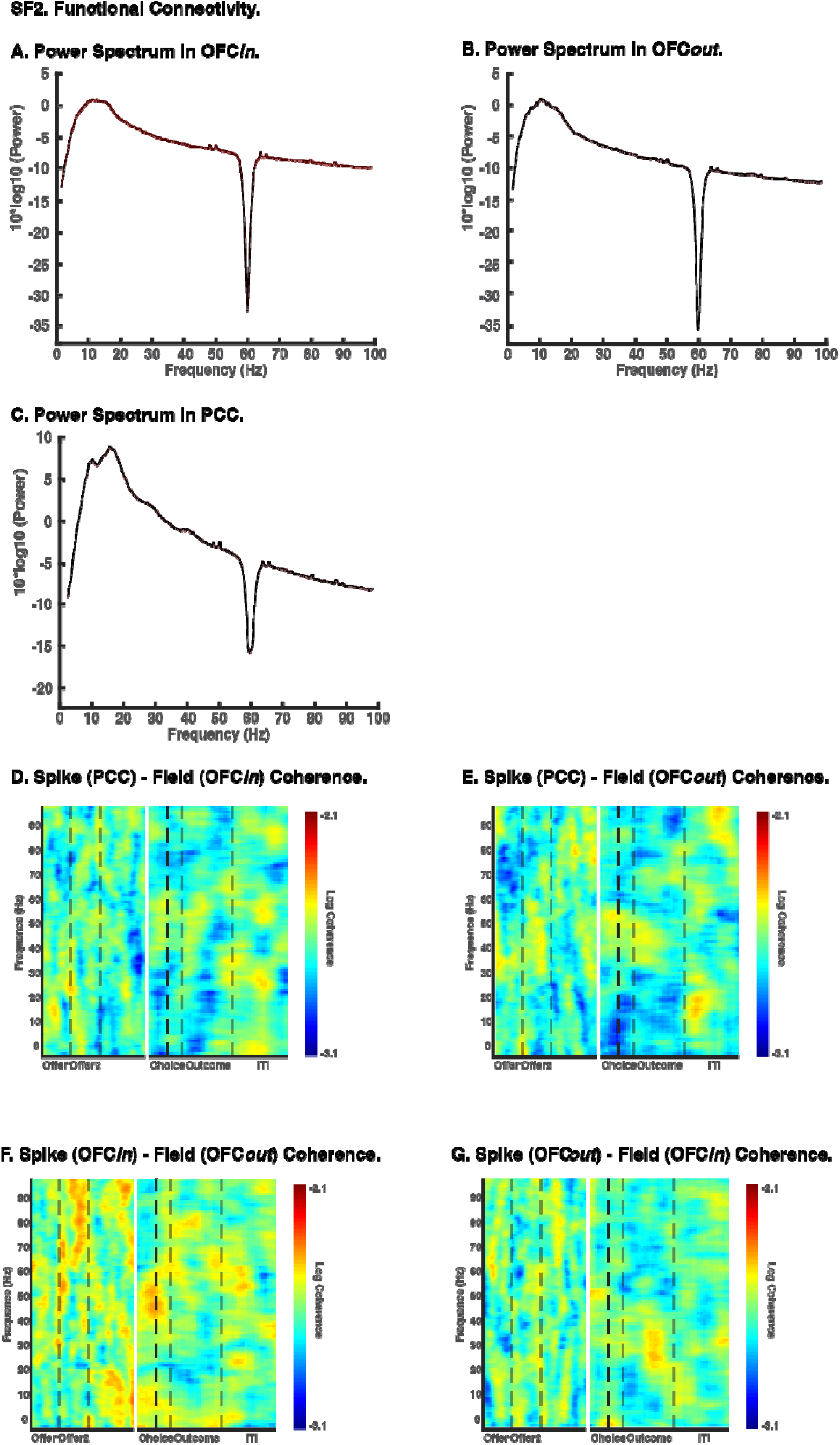
Supplementary Functional Connectivity. **A-C.** Power spectrum in OFC*in* (A), in OFC*out* (B), and in PCC (C). X axis: frequency (Hz). Y axis: power transformed with a 10 } *log10* function. Black line: mean power across channel and across trials. Red shaded area: 95% confidence interval. **D-J.** Spike-field coherence. X axis: time in a trial. Y axis: frequency. Color: strength of spike-field coherence on log10 scale. The warmer the color, the higher the coherence. Data from the first half of the trial (offer period) was aligned at offer 1 onset. Data from the second half of the trial (choice period) was aligned at Choice execution. (D) PCC_spk_-OFC*in*_lfp_ coherence. (E) PCC_spk_-OFC*out*_lfp_ coherence. (F) OFC*in*_spk_- OFC*out*_lfp_ coherence. (G) OFC*out*_spk_-OFC*in*_lfp_ coherence.

### Greater mutual information between OFC*in*-PCC and OFC*out*-PCC circuits

We found that the OFC*in*-PCC circuit shared more mutual information than OFC*out*-PCC (z=17.47, p<0.001, Wilcoxon signed rank test). Specifically, the OFC*in*-PCC circuit shared 7.44×10^−4^ bits of information per channel, while the OFC*out*-PCC circuit shared 6.72×10^−4^ bits per channel. This difference was observed during the offer 1 epoch (z=8.81, p<0.001), during the offer 2 epoch (z=8.34, p<0.001), during the choice epoch (z=9.42, p<0.001), and during the reward epoch (z=8.23, p<0.001). The difference was not observed during the inter-trial interval epoch (ITI, z=0.71, p=0.479).

### Encoding of offer, choice, and outcome

We next examined neural encoding of task parameters and behavior in OFC*in*, OFC*out*, and PCC using the proportion of neurons, the encoding strength, and the latency to peak encoding strength (**Methods**). All three regions encoded offer and outcome values with similar proportion of neurons, encoding strength, and latencies.

During the presentation of the first offer, 18.18% (n=8/44, p=0.001, binomial test) of OFC*in* neurons, 16.67% (n=9/54, p=0.001) of OFC*out* neurons, and 13.62% (n=29/213, p<0.001) of PCC neurons encoded the value of offer 1. These proportions were not detectably different from one another (χ^2^=0.79, df=2, p=0.675, Chi-square test).

We used the t-statistics of each predictor in a multiple regression model as a measure of encoding strength (**Methods**). Encoding strength of offer 1 value at the population level was not different among OFC*in*, OFC*out*, and PCC (χ^2^=1.67, p=0.434, Kruskal-Wallis test). We then assessed response latencies using a generalized linear model with Gamma distribution (Bishop, 2006; MacKay, 2003). For the latency analysis, we used all neurons, because many neurons in a population can show encoding of task variables without passing statistical significance; considering all neurons improves accuracy. Among all neurons, the encoding strength of offer 1 value peaked at 290 ms in OFC*in*, 235 ms in OFC*out*, and 240 ms in PCC, after offer 1 onset. We then used the distributions of single-neuron latencies to assess statistical significance; by this method, these latencies were not significantly different from one another (F=1.39, p=0.251, GLM with Gamma distribution; **Methods**).

During the outcome epoch, 34.09% (n=15/44, p<0.001, binomial test) of OFC*in* neurons, 35.19% (n=19/54, p<0.001, binomial test) of OFC*out* neurons, and 52.58% (n=112/213, p<0.001, binomial test) of PCC neurons encoded the value of received outcome. The proportion of such neurons in PCC was significantly higher than those of OFC*in* and OFC*out* (χ^2^=8.63, df=2, p=0.013, Chi-square test, cf. Hayden et al., 2008). The encoding strength of outcome value at the population level was significantly higher in PCC than both OFC*in* and OFC*out* (χ^2^=9.83, p=0.007, Kruskal-Wallis test with Tukey-Kramer multiple comparison). The encoding of outcome value peaked around 275 ms in OFC*in*, 360 ms in OFC*out*, and 450 ms in PCC, after reward onset. These latencies were not significantly different from one another (F=1.30, p=0.275, GLM with Gamma distribution).

All three regions encoded chosen option (offer 1 vs. 2) and chosen location (left vs. right). However, OFC*in* encoded the chosen option with shorter latency than both OFC*out* and PCC. PCC, not OFC*in* nor OFC*out*, showed a higher proportion of neurons encoding chosen location than chosen option. PCC and OFC*in* also encoded the chosen location with significantly shorter latencies than OFC*out*.

We defined choice epoch as the period from 200 ms after offer 2 was presented until when choice was made via saccade and fixation on the chosen option. During this time, 18.18% (n=8/44, p=0.001, binomial test) of OFC*in* neurons, 16.67% (n=9/54, p=0.001, binomial test) of OFC*out* neurons, and 12.21% (n=26/213, p=0.001, binomial test) of PCC neurons encoded chosen option (offer 1 vs. 2). These proportions were not significantly different from one another (χ^2^=2.62, df=2, p=0.270, Chi-square test). Encoding strength of chosen option at population level was not significantly different across regions (χ^2^=1.35, p=0.510, Kruskal-Wallis test). The encoding of chosen option peaked at 90 ms in OFC*in*, 170 ms in OFC*out*, and 150 ms in PCC into choice epoch. These latencies were significantly different from one another (F=3.35, p=0.037, GLM with Gamma distribution). Specifically, OFC*in* latency was significantly shorter than that in OFC*out* (t=−2.14, p=0.033, from the same GLM fit) or PCC (t=−2.36, p=0.019, from the same GLM fit), but there was no significant difference between OFC*out* and PCC (t=0.12, p=0.906, from the same GLM fit).

During the same choice epoch, 18.18% (n=8/44, p=0.001, binomial test) of OFC*in* neurons, 12.96% (n=7/54, p=0.018, binomial test) of OFC*out* neurons, and 19.25% (n=41/213, p<0.001, binomial test) of PCC neurons encoded chosen location (left vs. right). These proportions were not significantly different from one another (χ^2^=1.15, df=2, p=0.562, Chi-square test). However, PCC (χ^2^=5.31, df=1, p=0.021, Chi-square test) but not OFC*in* (χ^2^=0, df=1, p=1, Chi-square test) or OFC*out* (χ^2^=0.07, df=1, p=0.787, Chi-square test) showed a higher proportion of neurons encoding chosen location than chosen option. Encoding strength of chosen location at the population level was not significantly different across the three regions (χ^2^=0.20, p=0.906, Kruskal-Wallis test). The encoding of chosen location peaked around 150 ms in OFC*in*, 230 ms in OFC*out*, and 140 ms in PCC, into the choice epoch. These latencies were significantly different from one another (F=5.71, p=0.004, GLM with Gamma distribution). Specifically, OFC*out* latency was significantly longer than that in OFC*in* (t=2.36, p=0.019, from the same GLM fit) and PCC (t=3.47, p<0.001, from the same GLM fit), but there was no significant difference between those in OFC*in* and PCC (t=0.07, p=0.944, from the same GLM fit)‥

### Functional differences in decoding between OFC*in*-PCC and OFC*out*-PCC circuits

We then asked whether the relay of choice signal from value space to action space can be observed in decodability from population activities across all three regions. To answer this question, we took the normalized firing rate of each neuron over a sliding window to get the population activity pattern from all simultaneously recorded neurons in each trial. Then we trained a linear discriminant analysis (LDA) decoder on the population activity patterns from 75% of the trials and tested the decoder on the remaining 25% of the trials following a four-fold cross-validation procedure (**Methods**).

We found that at the end of offer 2 presentation (500 ms epoch), the value of the chosen option (offer 1 vs. 2) was decodable in all three of OFC*in* (χ^2^=7.41, p=0.006, chi-square test), OFC*out* (χ^2^=5.63, p=0.018), and PCC (χ^2^=12.45, p<0.001), on correct trials. These three proportions were not significantly different from one another (χ^2^=1.41, p=0.494), suggesting the decodability was similar across regions. The value of the chosen options was not decodable on error trials in OFC*in* (χ^2^=0.57, p=0.448) or PCC (χ^2^=0.25, p=0.613), although it was decodable in OFC*out* (χ^2^=6.83, p=0.009) (**Supplementary Figure 5A**). Right before a saccade was used to select the chosen option, chosen location (left vs. right) was not decodable in OFC*out* (χ^2^=0.02, p=0.901), but was decodable in OFC*in* (χ^2^=0.25, p=0.049) and PCC (χ^2^=8.85, p=0.003,), on correct trials. These three proportions were significantly different from one another (χ^2^=8.37, p=0.015); the proportion in PCC was significantly higher than in OFC*in* (χ^2^=8.12, p=0.004). Chosen location was not decodable on error trials in OFC*in* (χ^2^=0.30, p=0.584, OFC*out* (χ^2^=0, p=1), or PCC (χ^2^=0.06, p=0.801; **Supplementary Figure 5B**).

As a control test, we also tested decoding accuracy of EV1 high vs. low value. As shown in (**Supplementary Figure 5C-D**), both circuits / all three regions showed slightly but significantly higher than chance levels of decoding accuracy for whether EV1 was high or low in correct trials. Interestingly, decoding accuracies were not significantly different from the chance level in error trials during the offer period, and only reached slightly higher than chance level during outcome delivery.

Decodability for chosen location (left vs. right) was particularly prominent in PCC and was quenched on error trials. In addition, the decodability for chosen option (offer 1 vs. 2) in OFC*in* was also quenched on error trials. We wondered whether information in OFC*in* for chosen option, after being read out (decoded) by PCC, would influence the decodability of chosen location in PCC. In other words, does the relay of choice in value space to that in action space happen through the information input from OFC*in* to PCC, and can it be read out from PCC? To answer this question, we applied the Granger causality test to the decodability of chosen option and chosen location across regions.

We found that the decodability for chosen option (offer 1 vs. offer 2) in OFC*in* Granger-caused the decodability for chosen location (left vs. right) in PCC (gc=11.19, p=0.025) with a 200 ms (5 Hz) lag. This Granger-causal relation was absent on error trials (gc=3.04, p=0.552), suggesting the successful transform of choice signal in these two frameworks might be crucial for correct choice behavior. In the reverse direction, the decodability for chosen location (left vs. right) in PCC Granger-caused decodability for chosen option (offer 1 vs. 2) in OFC*in* (gc=17.59, p=0.025) but with a much longer time lag (400 ms; 2.5 Hz). In contrast, the decodability for chosen option (offer 1 vs. 2) in OFC*out* did not Granger-cause the decodability for chosen location (left vs. right) in PCC at any time lag, nor did the chosen location (left vs. right) in OFC*in* at any time lag.

### Functional differences in population dynamics between OFC*in*-PCC and OFC*out*-PCC circuits

After projecting trial-by-trial population states onto the top-N PC space, we found that in error trials, the overall distance between trial-by-trial population trajectories for chosen option (offer 1vs2) was significantly different across OFC*in*, OFC*out*, and PCC (χ^2^=59.88, p<0.001, Kruskal-Wallis test with Tukey-Kramer multiple comparison), with distance in OFC*out* significantly higher than in OFC*in* (p=0.036) and PCC (p<0.001) and distance in OFC*in* significantly higher than in PCC (p<0.001). The overall distance in error trials between trial-by-trial population trajectories for chosen location (left vs right) was not significantly different across OFC*in*, OFC*out*, and PCC (χ^2^=3.95, p=0.139). The overall distance between trial-by-trial population trajectories for high vs. low EV1 were significantly different across OFC*in*, OFC*out*, and PCC (χ^2^=8.83, p=0.012), with distance in OFC*in* significantly higher than that in OFC*out* (p=0.033) and PCC (p=0.023) but with no significant difference between OFC*out* and PCC (p=0.990).

We also compared the adjusted distance for trial-by-trial population trajectories between correct and error trials for different pairs of task parameters. In OFC*in*, adjusted distances between population trajectories for chosen option (offer 1vs2; χ^2^=61.82, p<0.001), chosen location (left vs right; χ^2^=111.99, p<0.001), and EV1 (high vs low; χ^2^=120.63, p<0.001) were significantly larger in correct than in error trials. In OFC*out* adjusted distances between population trajectories for chosen option (offer 1vs2; χ^2^=29.37, p<0.001), chosen location (left vs right; χ^2^=117.80, p<0.001), and EV1 (high vs low; χ^2^=137.78, p<0.001) were significantly larger in correct than in error trials. Similarly, in PCC, adjusted distances between population trajectories for chosen option (offer 1vs offer 2; χ^2^=93.01, p<0.001), chosen location (left vs right; χ^2^=137.49, p<0.001), and EV1 (high vs low; χ^2^=149.19, p<0.001) were significantly larger in correct than in error trials.

Simultaneously, all three regions showed larger dispersion (within-condition distance; see **Methods**) in error than in correct trials (**Supplementary Figure 6**). In OFC*in*, dispersion between population trajectories for chosen option (offer 1vs2; χ^2^=149.13, p<0.001, Kruskal-Wallis test), chosen location (left vs right; χ^2^=149.25, p<0.001), and EV1 (high vs low; χ^2^=149.25, p<0.001) were significantly larger in error than in correct trials. In OFC*out*, dispersion between population trajectories for chosen option (offer 1vs2; χ^2^=149.25, p<0.001), chosen location (left vs right; χ^2^=149.25, p<0.001), and EV1 (high vs low; χ^2^=149.25, p<0.001) were significantly larger in error than in correct trials. Similarly, in PCC, dispersion between population trajectories for chosen option (offer 1vs2; χ^2^=149.25, p<0.001), chosen location (left vs right; χ^2^=149.25, p<0.001), and EV1 (high vs low; χ^2^=149.25, p<0.001) were significantly larger in error than in correct trials.

**Supplementary Figure 4.**
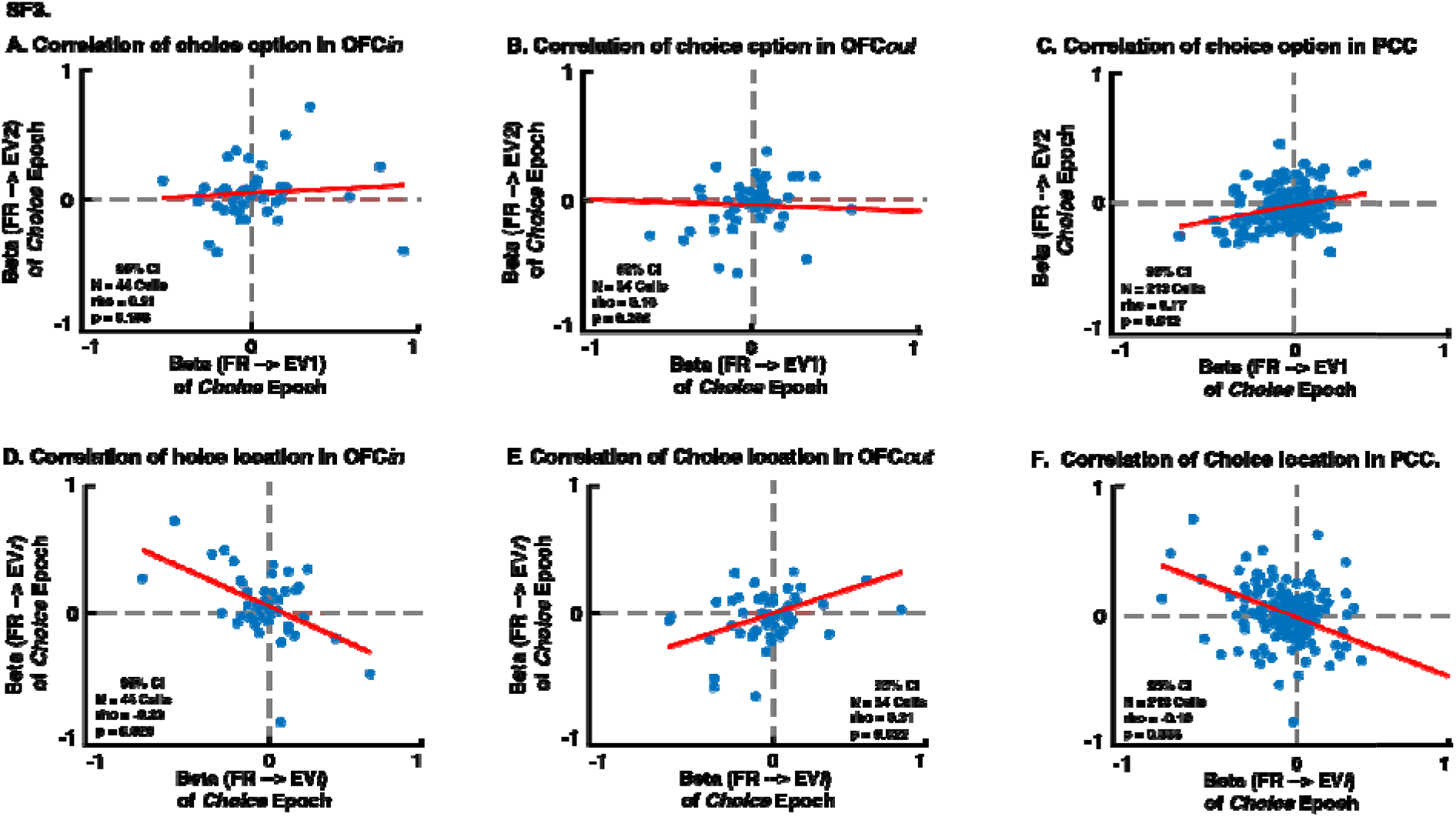
Putative mutual inhibition effects. **A-F**: Scatter plots. Each dot represents one neuron. Shaded area: 95% confidence interval. **A-C:** Y-axis: regression coefficient for expected value of offer 2. X-axis: regression coefficient for expected value of offer 1. **D-F:** Y-axis: regression coefficient for expected value of right offer. X-axis: regression coefficient for expected value of left offer. A,D: OFC*in*. B,E: OFC*out*. C,F: PCC. These figures are complementary to Figure 3A-F (main text) in that they are results from the same analysis in a later time window (from choice epoch instead of offer 2 epoch), to show the change and development of mutual inhibition signal.

**Supplementary Figure 5:**
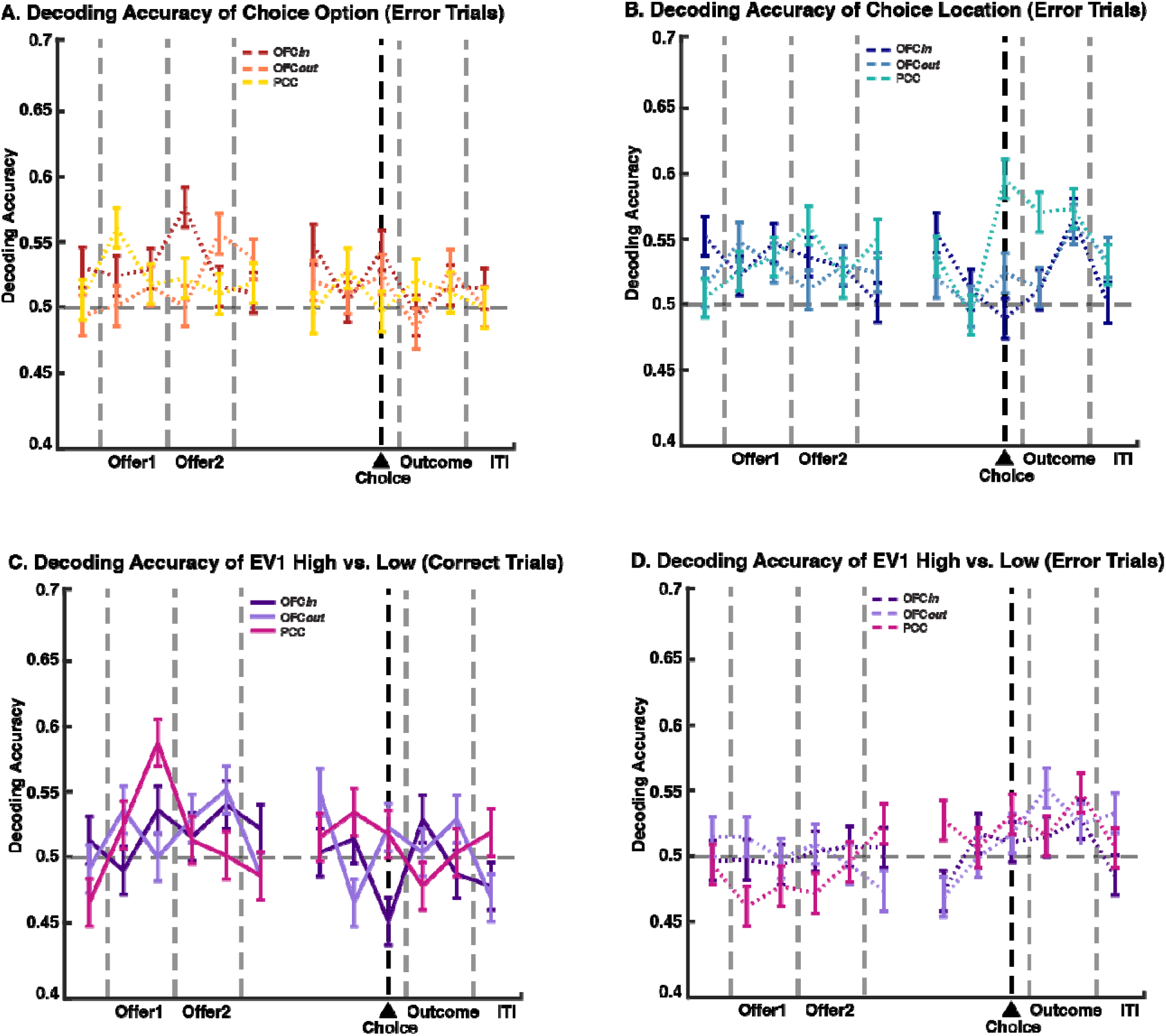
Decoding accuracy. **A-D**: Y-axis: probability of decoding correctly. X-axis: time in a trial. Error bar: standard error of the mean. **A:** Decoding accuracy of choice option (offer 1 vs. offer 2) from error trial (choosing the offer with the smaller expected value). **B:** Decoding accuracy of choice location (left vs. right) from error trials. **C-D**: Decoding accuracy of whether the expected value of offer 1 was higher or lower than the average expected value of offer 1 from correct (**C**) and error (**D**) trials, respectively.

**Supplementary Figure 6:**
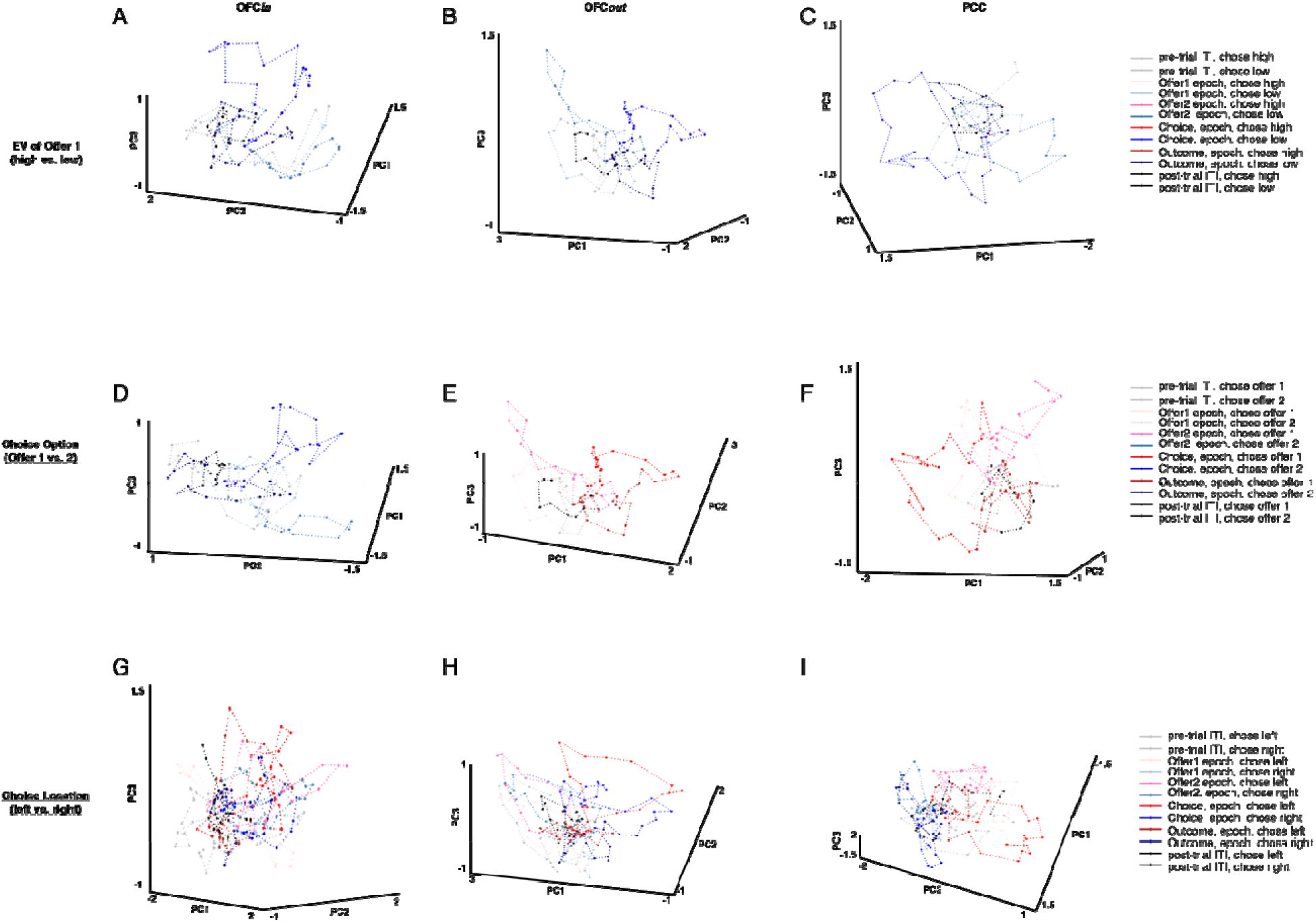
Population dynamics on error trials. Error trials are those in which the subject chose the offer with the smaller expected value. Trial averaged population activity projected onto top-N PC space (only top-3 PCs are shown here), separated by EV1 high vs. low (a-c), chosen option (offer 1 vs. offer 2; d-f), and chosen location (left vs. right; g-i), in OFCin (left column), OFCout (middle column), and PCC (right column). Warm color: trial averaged population activity for high EV1 (**A-C**), choosing offer 1 (**D-F**) or left offer (**G-I**). Cold color: trial averaged population activity for low EV1 (**A-C**), choosing offer 2 (**D-F**) or right offer (**G-I**). These figures are complementary to Figure 4. They show that the separate of trajectories based on EV1 and choices are diminished on error trials.

## REFERENCES

Afshar, A., Santhanam, G., Yu, B. M., Ryu, S. I., Sahani, M., & Shenoy, K. V. (2011). Single-Trial Neural Correlates of Arm Movement Preparation. Neuron, 71(3), 555–564.

Azab, H., & Hayden, B. Y. (2017). Correlates of decisional dynamics in the dorsal anterior cingulate cortex. PLoS biology, 15(11), e2003091.

Azab, H., & Hayden, B. Y. (2018). Correlates of economic decisions in the dorsal and subgenual anterior cingulate cortices. European Journal of Neuroscience, 47(8), 979–993.

Barack, D. L., Chang, S. W. C., & Platt, M. L. (2017). Posterior Cingulate Neurons Dynamically Signal Decisions to Disengage during Foraging. Neuron, 96(2), 339–347.e5.

Bartolo, R., & Averbeck, B. B. (2020). Prefrontal Cortex Predicts State Switches during Reversal Learning. Neuron, 1–11.

Bishop, C. M. (2006). Pattern Recoginiton and Machine Learning. In Information Science and Statistics.

Blanchard, T. C., & Hayden, B. Y. (2014). Neurons in dorsal anterior cingulate cortex signal postdecisional variables in a foraging task. Journal of Neuroscience, 34(2), 646–655.

Bokil, H., Andrews, P., Kulkarni, J. E., Mehta, S., & Mitra, P. P. (2010). Chronux: A platform for analyzing neural signals. Journal of Neuroscience Methods, 192(1), 146–151.

Bradfield, L. A., & Hart, G. (2020). Rodent medial and lateral orbitofrontal cortices represent unique components of cognitive maps of task space. Neuroscience and Biobehavioral Reviews, 108(June 2019), 287–294.

Brainard, D. H. (1997). The psychophysics toolbox. Spatial vision, 10(4), 433–436.

Buzsáki, G. (2004). Large-scale recording of neuronal ensembles. Nature Neuroscience, 7(5), 446–451.

Buzsaki, G., & Draguhn, A. (2004). Neuronal oscillations in cortical networks. Science, 304(5679), 1926–1929.

Cavada, C., & Goldman Rakic, P. S. (1989). Posterior parietal cortex in rhesus monkey: I. Parcellation of areas based on distinctive limbic and sensory corticocortical connections. Journal of Comparative Neurology, 287(4), 393–421.

Cai, X., & Padoa-Schioppa, C. (2014). Contributions of orbitofrontal and lateral prefrontal cortices to economic choice and the good-to-action transformation. Neuron, 81(5), 1140–1151.

Churchland, M. M., Cunningham, J. P., Kaufman, M. T., Foster, J. D., Nuyujukian, P., Ryu, S. I., Shenoy, K. V., & Shenoy, K. V. (2012). Neural population dynamics during reaching. Nature, 487(7405), 51–56.

Cornelissen, F. W., Peters, E. M., & Palmer, J. (2002). The Eyelink Toolbox: Eye tracking with MATLAB and the Psychophysics Toolbox. Behavior Research Methods, Instruments, and Computers.

Dal Monte, O., Chu, C. C. J., Fagan, N. A., & Chang, S. W. C. (2020). Specialized medial prefrontal–amygdala coordination in other-regarding decision preference. Nature Neuroscience.

Dean, H. L., Crowley, J. C., & Platt, M. L. (2004). Visual and saccade-related activity in macaque posterior cingulate cortex. Journal of neurophysiology, 92(5), 3056–3068.

Dean, H. L., & Platt, M. L. (2006). Allocentric spatial referencing of neuronal activity in macaque posterior cingulate cortex. Journal of Neuroscience.

Farashahi, S., Azab, H., Hayden, B., & Soltani, A. (2018). On the flexibility of basic risk attitudes in monkeys. Journal of Neuroscience, 38(18), 4383–4398.

Granger, C. W. J. (1969). Investigating Causal Relations by Econometric Models and Cross-Spectral Methods.”Econometrica. Vol. 37, 1969, pp. 424–459.

Haber, S. N., Kim, K. S., Mailly, P., & Calzavara, R. (2006). Reward-related cortical inputs define a large striatal region in primates that interface with associative cortical connections, providing a substrate for incentive-based learning. Journal of Neuroscience, 26(32), 8368–8376.

Hare, T. A., Schultz, W., Camerer, C. F., O’Doherty, J. P., & Rangel, A. (2011). Transformation of stimulus value signals into motor commands during simple choice. Proceedings of the National Academy of Sciences of the United States of America, 108(44), 18120–18125.

Hayden, B., Smith, D. V., & Platt, M. (2010). Cognitive control signals in posterior cingulate cortex. Frontiers in human neuroscience, 4, 223.

Hayden, B. Y., & Moreno-Bote, R. (2018). A neuronal theory of sequential economic choice. Brain and Neuroscience Advances.

Hayden, B. Y., Smith, D. V., & Platt, M. L. (2009). Electrophysiological correlates of default-mode processing in macaque posterior cingulate cortex. Proceedings of the National Academy of Sciences, 106(14), 5948–5953.

Hayden, B. Y., Nair, A. C., McCoy, A. N., & Platt, M. L. (2008). Posterior Cingulate Cortex Mediates Outcome-Contingent Allocation of Behavior. Neuron, 60(1), 19–25.

Hayden, B. Y., & Heilbronner, S. R. (2014). All that glitters is not reward signal. Nature neuroscience, 17(9), 1142–1144.

Heilbronner, S. R., & Platt, M. L. (2013). Causal evidence of performance monitoring by neurons in posterior cingulate cortex during learning. Neuron, 80(6), 1384–1391.

Heilbronner, S. R., & Hayden, B. Y. (2016). Dorsal anterior cingulate cortex: a bottom-up view. Annual review of neuroscience, 39, 149–170.

Heilbronner, S. R. (2017). Modeling risky decision-making in nonhuman animals: shared core features. Current opinion in behavioral sciences, 16, 23–29.

Heilbronner, S. R., & Hayden, B. Y. (2016). The description-experience gap in risky choice in nonhuman primates. Psychonomic bulletin & review, 23(2), 593–600.

Kable, J. W., & Glimcher, P. W. (2007). The neural correlates of subjective value during intertemporal choice. Nature Neuroscience, 10(12), 1625–1633.

Kaplan, R., Schuck, N. W., & Doeller, C. F. (2017). The Role of Mental Maps in Decision-Making. Trends in Neurosciences, 40(5), 256–259.

Kobayashi, Y., & Amaral, D. G. (2003). Macaque monkey retrosplenial cortex: II. Cortical afferents. Journal of Comparative Neurology, 466(1), 48–79.

Leech, R., & Sharp, D. J. (2014). The role of the posterior cingulate cortex in cognition and disease. Brain, 137(1), 12–32.

Levy, D. J., & Glimcher, P. W. (2012). The root of all value: a neural common currency for choice. Current opinion in neurobiology, 22(6), 1027–1038.

Luk, C. H., & Wallis, J. D. (2013). Choice coding in frontal cortex during stimulus-guided or action-guided decision-making. Journal of Neuroscience, 33(5), 1864–1871.

Lütkepohl, H. (2007). New Introduction to Multiple Time Series Analysis. New York, NY: Springer-Verlag.

MacKay, D. J. C. (2003). Information Theory, Inference and Learning Algorithms. Cambridge University Press.

Mante, V., Sussillo, D., Shenoy, K. V., & Newsome, W. T. (2013). Context-dependent computation by recurrent dynamics in prefrontal cortex. Nature, 503(7474), 78–84.

Morecraft, R. J., Cipolloni, P. B., Stilwell-Morecraft, K. S., Gedney, M. T., & Pandya, D. N. (2004). Cytoarchitecture and Cortical Connections of the Posterior Cingulate and Adjacent Somatosensory Fields in the Rhesus Monkey. Journal of Comparative Neurology.

Morecraft, R. J., Geula, C., & Mesulam, M. M. (1992). Cytoarchitecture and neural afferents of orbitofrontal cortex in the brain of the monkey. Journal of Comparative Neurology, 323(3), 341–358.

Mufson, E. J., & Pandya, D. N. (1984). Some observations on the course and composition of the cingulum bundle in the rhesus monkey. Journal of Comparative Neurology, 225(1), 31–43.

Murray, J. D., Bernacchia, A., Roy, N. A., Constantinidis, C., Romo, R., & Wang, X. J. (2017). Stable population coding for working memory coexists with heterogeneous neural dynamics in prefrontal cortex. Proceedings of the National Academy of Sciences of the United States of America, 114(2), 394–399.

Murray, E. A., & Rudebeck, P. H. (2018). Specializations for reward-guided decision-making in the primate ventral prefrontal cortex. Nature Reviews Neuroscience, 19(7), 404–417.

Niv, Y. (2019). Learning task-state representations. Nature neuroscience, 22(10), 1544–1553.

Noonan, M. P., Walton, M. E., Behrens, T. E. J., Sallet, J., Buckley, M. J., & Rushworth, M. F. S. (2010). Separate value comparison and learning mechanisms in macaque medial and lateral orbitofrontal cortex. Proceedings of the National Academy of Sciences, 107(47), 20547–20552.

O’Doherty, J. P. (2014). The problem with value. Neuroscience & Biobehavioral Reviews, 43, 259–268.

Olson, C. R., Musil, S. Y., & Goldberg, M. E. (1996). Single neurons in posterior cingulate cortex of behaving macaque: Eye movement signals. Journal of Neurophysiology, 76(5), 3285–3300.

Öngür, D., & Price, J. L. (2000). The organization of networks within the orbital and medial prefrontal cortex of rats, monkeys and humans. In Cerebral Cortex.

Padoa-Schioppa, C. (2011). Neurobiology of Economic Choice: A Good-Based Model. Annual Review of Neuroscience, 34(1), 333–359.

Padoa-Schioppa, C., & Conen, K. E. (2017). Orbitofrontal Cortex: A Neural Circuit for Economic Decisions. Neuron, 96(4), 736–754.

Pandya, D. N., Van Hoesen, G. W., & Mesulam, M. M. (1981). Efferent connections of the cingulate gyrus in the rhesus monkey. Experimental Brain Research.

Pandya, D. N., & Seltzer, B. (1982). Intrinsic connections and architectonics of posterior parietal cortex in the rhesus monkey. Journal of Comparative Neurology, 204(2), 196–210.

Parvizi, J., Van Hoesen, G. W., Buckwalter, J., & Damasio, A. (2006). Neural connections of the posteromedial cortex in the macaque. Proceedings of the National Academy of Sciences of the United States of America.

Paxinos G, Huang XF, Petrides M, T. A. (2009). The Rhesus Monkey Brain in Stereotaxic Coordinates. Academic Press.

Pearson, J. M., Hayden, B. Y., Raghavachari, S., & Platt, M. L. (2009). Neurons in Posterior Cingulate Cortex Signal Exploratory Decisions in a Dynamic Multioption Choice Task. Current Biology.

Pesaran, B. (2010). Neural correlations, decisions, and actions. Current Opinion in Neuobiology, 20(2), 166–171.

Pirrone, A., Azab, H., Hayden, B. Y., Stafford, T., & Marshall, J. A. (2018). Evidence for the speed–value trade-off: Human and monkey decision making is magnitude sensitive. Decision, 5(2), 129.

Rangel, A., Camerer, C., & Montague, P. R. (2008). A framework for studying the neurobiology of value-based decision making. Nature reviews neuroscience, 9(7), 545–556.

Roesch, M. R., Taylor, A. R., & Schoenbaum, G. (2006). Encoding of time-discounted rewards in orbitofrontal cortex is independent of value representation. Neuron, 51(4), 509–520.

Rudebeck, P. H., & Murray, E. A. (2011). Balkanizing the primate orbitofrontal cortex: distinct subregions for comparing and contrasting values. Annals of the New York Academy of Sciences, 1239, 1.

Rudebeck, P. H., Saunders, R. C., Lundgren, D. A., & Murray, E. A. (2017). Specialized representations of value in the orbital and ventrolateral prefrontal cortex: desirability versus availability of outcomes. Neuron, 95(5), 1208–1220.

Rushworth, M. F. S., Noonan, M. P., Boorman, E. D., Walton, M. E., & Behrens, T. E. (2011). Frontal cortex and reward-guided learning and decision-making. Neuron, 70(6), 1054–1069.

Scherberger, H., Jarvis, M. R., & Andersen, R. A. (2005). Cortical Local Field Potential Encodes Movement Intentions in the Posterior Parietal Cortex. Neuron, 46(2), 347–354.

Schoenbaum, G., Roesch, M. R., Stalnaker, T. A., & Takahashi, Y. K. (2009). A new perspective on the role of the orbitofrontal cortex in adaptive behaviour. Nature Reviews Neuroscience, 10(12), 885–892.

Schuck, N. W., Cai, M. B., Wilson, R. C., & Niv, Y. (2016). Human Orbitofrontal Cortex Represents a Cognitive Map of State Space. Neuron, 91(6), 1402–1412.

Sleezer, B. J., Castagno, M. D., & Hayden, B. Y. (2016). Rule encoding in orbitofrontal cortex and striatum guides selection. Journal of Neuroscience, 36(44), 11223–11237.

Spreng, R. N., Mar, R. A., & Kim, A. S. N. (2009). The common neural basis of autobiographical memory, prospection, navigation, theory of mind, and the default mode: A quantitative meta-analysis. Journal of Cognitive Neuroscience, 21(3), 489–510.

Stalnaker, T. A., Cooch, N. K., & Schoenbaum, G. (2015). What the orbitofrontal cortex does not do. Nature Neuroscience, 18(5), 620.

Strait, C. E., Blanchard, T. C., & Hayden, B. Y. (2014). Reward value comparison via mutual inhibition in ventromedial prefrontal cortex. Neuron, 82(6), 1357–1366.

Strait, C. E., Sleezer, B. J., & Hayden, B. Y. (2015). Signatures of value comparison in ventral striatum neurons. PLoS Biol, 13(6), e1002173.

Strait, C. E., Sleezer, B. J., Blanchard, T. C., Azab, H., Castagno, M. D., & Hayden, B. Y. (2016). Neuronal selectivity for spatial positions of offers and choices in five reward regions. Journal of neurophysiology, 115(3), 1098–1111.

Timme, N. M., & Lapish, C. (2018). A tutorial for information theory in neuroscience. ENeuro, 5(3).

Vickery, T. J., Chun, M. M., & Lee, D. (2011). Ubiquity and specificity of reinforcement signals throughout the human brain. Neuron, 72(1), 166–177

Vogt, B. A., & Paxinos, G. (2014). Cytoarchitecture of mouse and rat cingulate cortex with human homologies. Brain Structure and Function.

Wallis, J. D. (2007). Orbitofrontal cortex and its contribution to decision-making. Annual Review of Neuroscience, 30(1), 31–56.

Wang, F., Schoenbaum, G., & Kahnt, T. (2020). Interactions between human orbitofrontal cortex and hippocampus support model-based inference. PLoS Biology, 18(1), e3000578.

Wang, M. Z., & Hayden, B. Y. (2017). Reactivation of associative structure specific outcome responses during prospective evaluation in reward-based choices. Nature Communications, 8(May), 1–13.

Widge, A. S., Heilbronner, S. R., & Hayden, B. Y. (2019). Prefrontal cortex and cognitive control: new insights from human electrophysiology. F1000Research, 8.

Wikenheiser, A. M., & Schoenbaum, G. (2016). Over the river, through the woods: Cognitive maps in the hippocampus and orbitofrontal cortex. Nature Reviews Neuroscience, 17(8), 513–523.

Wilson, R. C., Takahashi, Y. K., Schoenbaum, G., & Niv, Y. (2014). Orbitofrontal cortex as a cognitive map of task space. Neuron, 81(2), 267–279.

Yim, M. Y., Cai, X., & Wang, X. J. (2019). Transforming the Choice Outcome to an Action Plan in Monkey Lateral Prefrontal Cortex: A Neural Circuit Model. Neuron, 103(3), 520–532.e5.

Yoo, S. B. M., Sleezer, B. J., & Hayden, B. Y. (2018). Robust encoding of spatial information in orbitofrontal cortex and striatum. Journal of Cognitive Neuroscience, 30(6), 898–913.

Yoo, S. B. M., & Hayden, B. Y. (2020). The transition from evaluation to selection involves neural subspace reorganization in core reward regions. Neuron, 105(4), 712–724.

Yoo, S. B. M., & Hayden, B. Y. (2018). Economic choice as an untangling of options into actions. Neuron, 99(3), 434–447.

Young, M. E., & Mccoy, A. W. (2015). A delay discounting task produces a greater likelihood of waiting than a deferred gratification task. Journal of the Experimental Analysis of Behavior, 103(1), 180–195.

